# Statistical significance testing for spatial domain detection in spatial transcriptomics

**DOI:** 10.64898/2026.11.07.765075

**Authors:** Huairui Wang, Feiyu Cao, Hui Shen, Yufeng Liu, Guanao Yan

## Abstract

In spatial transcriptomics, domain-detection methods group platform-specific spatial measurement units—cells, bins or multicellular spots—into spatial domains for comparing molecular programs, cellular composition and interactions among tissue regions. However, the number of candidate domains detected in the same tissue can vary with user-specified parameters or model-selection criteria. Whether candidate domains should be merged can be framed as a test of a single-domain null hypothesis, under which each gene has constant mean expression across all spatial units in the combined region. However, testing candidate domains on the same gene-expression data used to detect them may induce post-selection bias and yield anti-conservative *p*-values, increasing the risk of oversegmentation. To address these challenges, we introduce SigDoTS (statistical Significance testing for Domain deTection in Spatial transcriptomics), a hypothesis-testing framework for spatial domain detection. SigDoTS uses hierarchical testing to determine which candidate domains to retain or merge, with adjustment for family-wise error control. It accounts for gene–gene covariance and spatial dependence and computes *p*-values by rerunning the upstream domain-detection pipeline on datasets simulated under the fitted single-domain null. In simulations, SigDoTS controlled the type I error rate at the root node under the single-domain null and improved recovery of the true spatial domains when multiple domains were present. In human dorsolateral prefrontal cortex, SigDoTS retained superficial and deep Layer 6 domains but merged Layer 3 candidate domains across different initial numbers of candidate domains. In ovarian and colorectal cancers, SigDoTS retained epithelial domains with different gene-expression programs and local immune environments, together with immune, stromal and vascular domains.

## Introduction

In spatial transcriptomics, domain-detection methods group platform-specific spatial measurement units—cells, bins or multicellular spots—into spatial domains [1–5]. The resulting domains are commonly described as spatially coherent tissue regions with similar gene-expression profiles [3, 4, 6] and provide the region labels used in downstream analyses [7, 8], such as spatially variable gene detection [4, 9, 10] and cell–cell interaction analyses. However, the number of domains returned by a domain-detection method depends on its input or tuning criterion. Depending on the method, this number is specified directly, induced by a clustering-resolution parameter or selected using a model-selection criterion [1, 2, 11–13]. Different choices can return different numbers of spatial domains for the same tissue [14], and thus different regions for downstream analyses, yet how to choose this number remains unclear.

To address this, we formulate the question of whether a tested region should be divided as a statistical hypothesis test and define the single-domain null hypothesis: each gene has constant mean expression across all spatial units in the tested region, although expression can be correlated across genes and across neighboring spatial units. We define oversegmentation as retaining multiple spatial domains within a region that satisfies the single-domain null. For example, a domain-detection method set to return two domains does so even when every gene has the same mean expression across all spatial units in the entire analyzed region (the global null) [15–17] (Fig. 1a).

**Figure 1:**
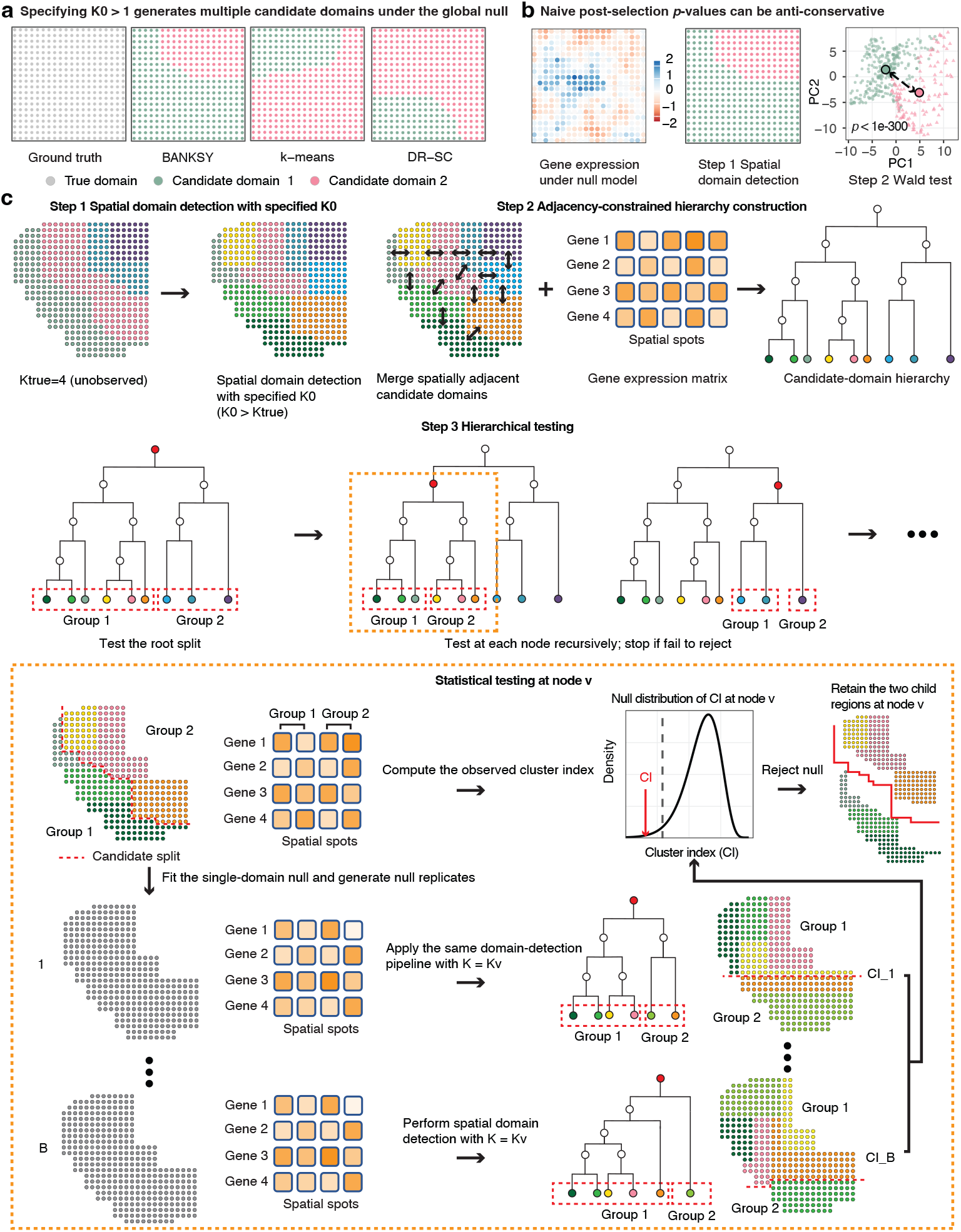
Overview of the motivation and workflow of SigDoTS. **a** When the number of initial candidate domains, *K*_0_>, is set greater than one, a domain-detection method divides data generated under the global null into multiple candidate domains. **b** Treating candidate domains detected from the gene-expression data as prespecified in subsequent testing can yield anti-conservative *p*-values. **c** SigDoTS proceeds in three steps. At node *v, K*_*v*_ denotes the number of initial candidate domains descending from that node. The inset illustrates the nodewise two-domain test.

Testing spatial domains introduces an additional inferential challenge: the same gene-expression data are used to detect spatial domains and to test their mean-expression separation (“double dipping”); a test that treats these domains as prespecified is therefore subject to post-selection bias [9, 18] (Fig. 1b). This bias can yield anti-conservative *p*-values and increase the probability of rejecting a true single-domain null [19–21]. Thus, oversegmentation can arise when the domains returned by a domain-detection method are retained without testing, or when they are tested as if prespecified. Two lines of hypothesis-testing work address analogous problems in non-spatial data. Simulation-based tests compare the observed clusters with clusters obtained by applying the same clustering procedure to data generated under a single-population null, as in SigClust [15], its hierarchical extension SHC, which tests each two-way split in a clustering hierarchy [16], and the single-cell adaptation scSHC [17]. Selective inference instead tests the difference in means between two estimated clusters conditional on the clustering event [19, 20]. Both assume independent observations, whereas neighboring cells, bins or spots in spatial transcriptomics often have correlated profiles; null data generated under independence can then underestimate the separation that clustering produces, yielding anti-conservative tests. Selective inference has been extended to dependent data [22] but applies only to clustering algorithms for which the datasets yielding the same clusters can be described in closed form, such as hierarchical and *k*-means clustering, and not to most spatial domain-detection methods [3–5, 11, 12].

To address these challenges, we introduce SigDoTS (statistical Significance testing for Domain deTection in Spatial transcriptomics), a hypothesis-testing framework for limiting oversegmentation in spatial domain detection (Fig. 1c). SigDoTS arranges the *initial candidate domains* returned by a domain-detection method in a binary hierarchy based on gene-expression similarity and spatial adjacency, so that each test evaluates a *candidate split*: a proposed division of one tested region into two. It tests candidate splits from the root (the entire analyzed region) downward against a matrix-normal single-domain null that models both gene–gene covariance and spatial dependence. At each candidate split, it generates replicate datasets from the fitted single-domain null and reruns the same domain-detection method on each replicate. Each *p*-value therefore compares the observed separation with separations produced by the same method when the tested region follows the single-domain null. SigDoTS uses hierarchical adjustment to control the family-wise error rate. We evaluated SigDoTS in simulations with spatially correlated expression and applied it to human dorsolateral prefrontal cortex profiled by Visium, ovarian cancer profiled by Visium HD and colorectal adenocarcinoma profiled by Xenium.

## Results

### SigDoTS controls type I error and merges oversegmented candidate domains in simulations

We simulated gene count matrices for *n* spatial units on a square lattice and *p* genes from a Poisson log-normal model [23], using a Matérn spatial covariance to introduce correlation between nearby spatial units (Supplementary Methods S1.2). We compared SigDoTS with two baselines: scSHC [17] and a hierarchical-clustering Wald test (HC-Wald). HC-Wald tested the same hierarchy with the same hierarchical adjustment as SigDoTS (Methods) but treated candidate domains as prespecified, comparing the mean scores of the two child regions on the first 15 principal components of gene expression with a Wald test. As an untested reference, we also report the initial candidate domains returned by *k*-means (“Observed” in Fig. 2). We denote the true number of spatial domains by *K*_true_, the number of initial candidate domains returned by the domain-detection method by *K*_0_> and the number of domains retained after testing by *K*.

**Figure 2:**
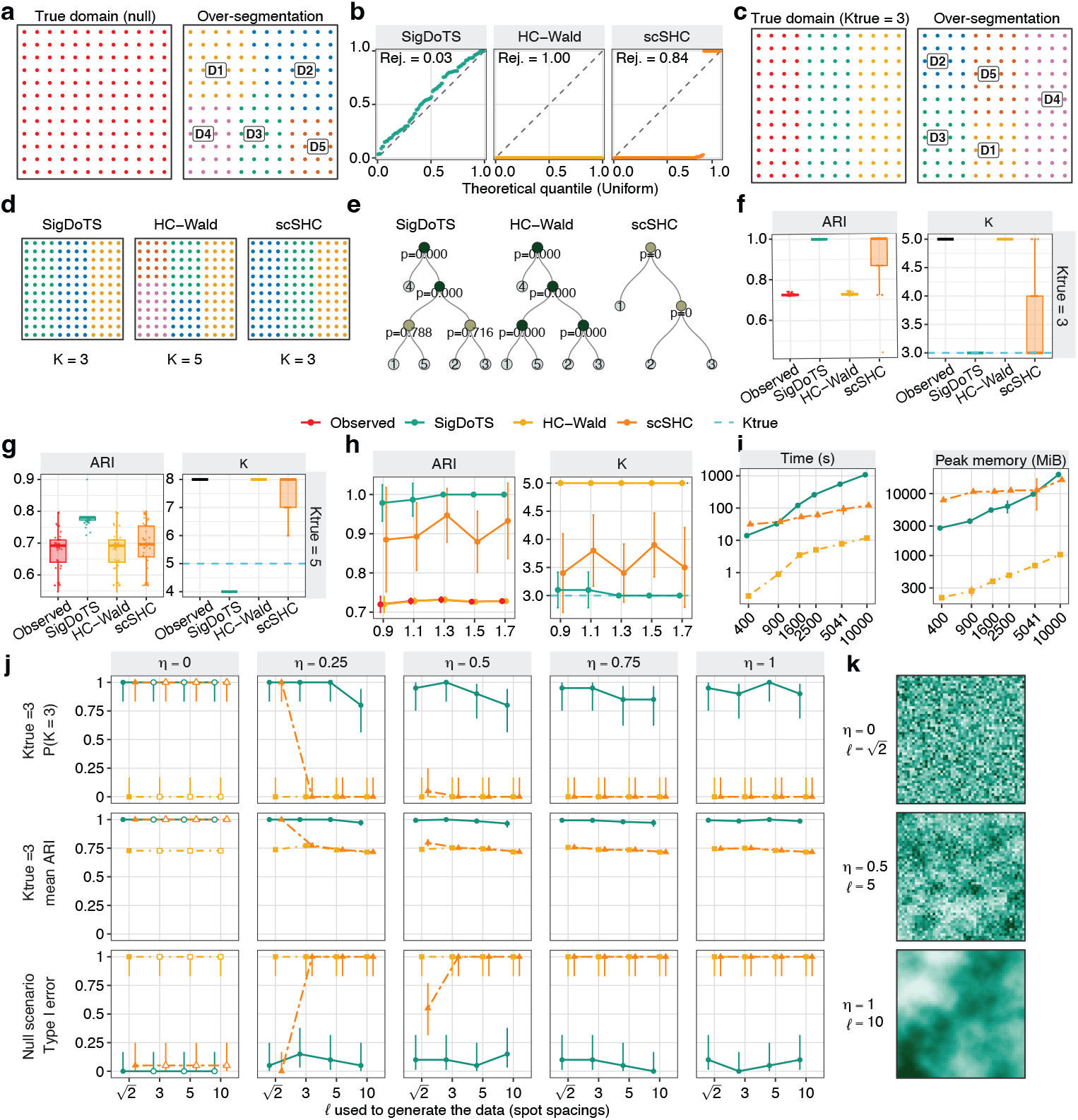
Spatial domain recovery and null calibration in Poisson log-normal simulations. **a** A global-null dataset (*K*_true_ = 1) and its *K*_0_> = 5 initial candidate domains. **b** Root *p*-value quantile–quantile plots for 100 global-null datasets; “Rej.” denotes the root rejection rate at *α* = 0.05. **c**–**e** A three-domain alternative dataset (*K*_true_ = 3, δ = 1.3): true domains and *K*_0_> = 5 initial candidate domains (**c**), retained domains (**d**) and testing trees (**e**). **f**,**g** Adjusted Rand index (ARI) and retained *K* across 40 three-domain (**f**; *K*_0_> = 5) and 40 five-domain (**g**; *K*_0_> = 8) alternative datasets; “Observed” denotes the initial candidate domains. **h** Mean ARI and retained *K* across signal strengths δ (10 datasets per setting). **i** Mean runtime and peak memory across 400–10,000 spatial units (two datasets per setting; 10 processor cores per method). **j** Proportion of three-domain datasets with *K* = *K*_true_, mean ARI and global-null root rejection rate across spatial covariance settings (*η, l*) on a 50 × 50 lattice (20 datasets per setting). For = 0, data were generated only at 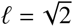 open symbols repeat these values. **k** Log-normalized expression of one gene in a global-null dataset at three (*η, l*) settings.

First, under the global null (*K*_true_ = 1), SigDoTS controlled the type I error, whereas both baselines retained multiple domains in at least 84% of datasets. In this scenario, every gene had the same mean expression across all spatial units, yet *k*-means with *K*_0_> = 5 divided each dataset into five initial candidate domains. We generated 100 datasets with 2,000 genes at each of two sizes, 144 and 2,000 spatial units. SigDoTS rejected the root null hypothesis in 3% and 1% of datasets at the two sizes (*α* = 0.05), merging the five initial candidate domains into a single domain in all other datasets, and its root *p*-values were approximately uniform (Fig. 2a,b; Supplementary Fig. S1a). HC-Wald, which treats candidate domains as prespecified, rejected the root null in every dataset at both sizes, as expected from double dipping. scSHC, which accounts for the clustering step but generates null data without spatial correlation, rejected it in 84% of datasets at 144 spatial units and in all datasets at 2,000.

Second, when the data contained three true spatial domains (*K*_true_ = 3), SigDoTS merged the oversegmented candidate domains and retained the three true domains. We shifted the latent log-rate mean of 200 randomly selected genes to − δ/2, +δ/2 and +2δ across the three domains (δ = 1.3); the remaining genes retained a mean of zero. *k*-means with *K*_0_> = 5 divided the three true domains into five initial candidate domains (mean adjusted Rand index (ARI), 0.729). SigDoTS retained exactly three domains in 39 of 40 datasets and increased the mean ARI to 0.997 (Fig. 2c–f; Supplementary Fig. S1b). HC-Wald retained all five initial candidate domains in every dataset, leaving the oversegmentation unchanged, whereas scSHC retained three domains in 26 of 40 datasets (mean ARI, 0.936; Fig. 2f).

Third, with five true domains (*K*_true_ = 5, *K*_0_> = 8), SigDoTS achieved the highest ARI of all methods, and its errors at the smaller sample size were in the conservative direction. At *n* = 400, no method recovered all five domains in any of the 40 datasets; SigDoTS retained four domains in every dataset, one fewer than the truth, and achieved the highest mean ARI (0.772, compared with 0.675 for HC-Wald and the initial candidate domains and 0.673 for scSHC; Fig. 2g). At *n* = 2,000, SigDoTS retained a median of five domains and again achieved a higher ARI than both baselines (Supplementary Fig. S1c), consistent with limited power rather than systematic merging at *n* = 400.

We then assessed the sensitivity of SigDoTS to the upstream domain-detection method, the initial candidate-domain count, signal strength, domain-size imbalance and the number of spatial units (Supplementary Results S2.1). With initial candidate domains from BANKSY [11], DR-SC [12] or *k*-means (*K*_0_> = 10), SigDoTS controlled the type I error and recovered the three true domains (Supplementary Fig. S2a–e). Across *K*_0_> ∈ {4, 6, 9, 12, 18}, it also controlled the type I error and increased the ARI relative to the initial candidate domains, although the ARI and boundary F1 score (Supplementary Methods S1.7) decreased at *K*_0_> = 18, the largest *K*_0_> tested (Supplementary Fig. S2f–h). Across signal strengths from δ = 0.9 to 1.7 in the three-domain alternative (*K*_true_ = 3, *K*_0_> = 5), SigDoTS reduced the oversegmentation at every setting; even at the weakest signal, it retained three domains in 9 of 10 datasets (mean ARI, 0.979; Fig. 2h, Supplementary Fig. S1d). When the first of three domains varied from half to three times the size of the other two, SigDoTS retained the three true domains in 37 to 40 of 40 datasets per setting when this domain was at least as large as the others and two domains in 36 of 40 datasets when it was half their size (Supplementary Fig. S2i–l); at 2,000 spatial units, it recovered all three domains at every tested imbalance (Supplementary Fig. S1e). As the number of spatial units increased from 400 to 10,000, SigDoTS recovered the three true domains exactly (ARI, 1.000) in both datasets at every size, whereas HC-Wald retained all five initial candidate domains and scSHC recovered three domains only at 10,000 spatial units. Over this 25-fold increase, the runtime of SigDoTS increased 77-fold, from 14 to 1,076 seconds, and its peak memory 7.3-fold, from 2,778 to 20,283 MiB (Fig. 2i).

Finally, as spatial correlation increased in strength or range, SigDoTS maintained domain recovery, whereas the global-null rejection rate of scSHC rose to 100%. We varied the strength, measured by the fraction of spatially correlated variance, and the range, measured by the length scale *l*, on a 50 × 50 lattice. Under the three-domain alternative, SigDoTS recovered the three true domains in 80% to 100% of datasets per setting and matched or exceeded the mean ARI of both baselines in every setting (Fig. 2j). Under the global null, SigDoTS rejected the root null in 0 to 3 of 20 datasets per setting (25 of 340 in total, 7.4%, compared with the nominal 5%), and HC-Wald rejected it in all datasets. scSHC controlled the type I error only when spatial units were uncorrelated ( = 0). As and *l* increased, expression varied more smoothly without domain-specific mean differences (Fig. 2k).

### SigDoTS separates superficial and deep Layer 6 and merges Layer 3 candidate domains in human dorsolateral prefrontal cortex

We applied SigDoTS at *α* = 0.05 to four human dorsolateral prefrontal cortex (DLPFC) Visium sections with manually annotated cortical layers (Supplementary Methods S1.3). First, we evaluated whether SigDoTS retained the reported within-layer organization in DLPFC Layer 6, where previous spatial and single-cell analyses identified molecularly distinct subdomains [14]. We included all manually labeled Layer 6 spots (Fig. 3a) from the four sections. BANKSY generated candidate domains at *K*_0_> = 3 (Fig. 3b). SigDoTS consistently retained two domains corresponding to superficial and deep Layer 6 across all four sections (root *p* = 0.008; Fig. 3c–d). In contrast, HC-Wald retained all candidate domains in all four sections (Supplementary Fig. S3a), and scSHC retained two, one, three and two domains across the four sections, respectively (Fig. 3c, Supplementary Fig. S3b); its boundaries varied among sections and did not form the same superficial–deep division. Across all four sections, superficial Layer 6 had higher mean standardized expression of neuronal markers, whereas deep Layer 6 had higher expression of oligodendrocyte/myelination markers (Fig. 3e). Along normalized cortical depth, the neuronal module score decreased, whereas the oligodendrocyte/myelination module score increased (Fig. 3f,g).

**Figure 3:**
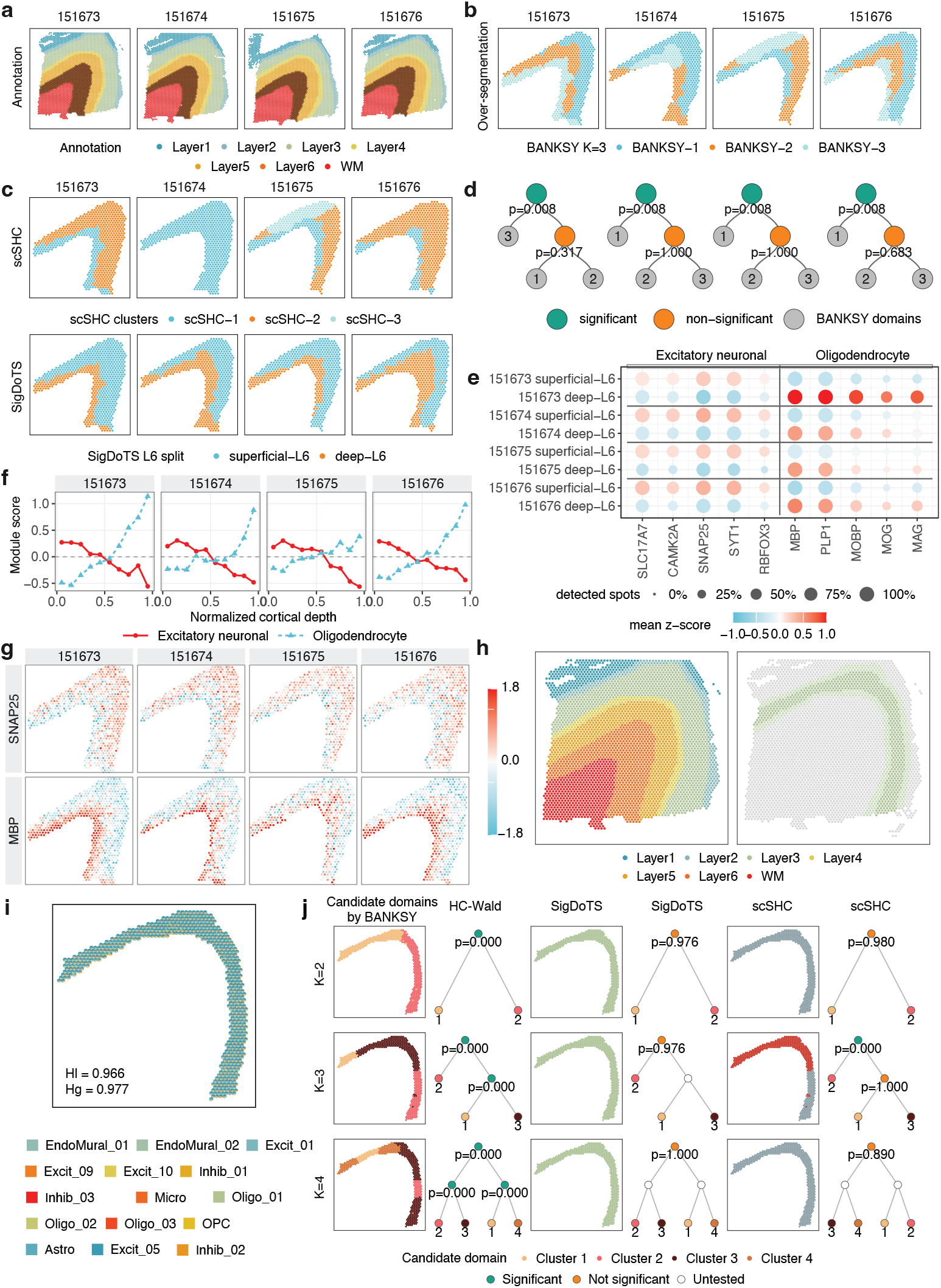
Layer 6 subdivision and Layer 3 interior testing in human dorsolateral prefrontal cortex. **a** Manual cortical-layer annotations. **b** Initial BANKSY candidate domains within Layer 6 (*K*_0_> = 3). **c** Domains retained by scSHC (top) and SigDoTS (bottom). **d** SigDoTS testing trees for Layer 6; node labels give *p*-values, green and orange nodes denote rejection and failure to reject, and gray leaves identify the initial candidate domains. **e** Marker expression in superficial and deep Layer 6 (color, mean standardized expression; dot size, percentage of spots expressing each gene). **f** Excitatory-neuronal and oligodendrocyte/myelination module scores along normalized cortical depth, from the Layer 5-facing edge (0) to the white-matter-facing edge (1). **g** Standardized *SNAP25* and *MBP* expression within Layer 6. **h** Section 151673 (left) and the selected Layer 3 interior (right). **i** Cell2location-estimated cell-type proportions within the Layer 3 interior (spot-level pie charts), with local (*H*_*l*_) and global (*H*_*g*_) composition homogeneity. **j** BANKSY candidate domains, testing trees and retained domains for the Layer 3 interior at *K*_0_> = 2, 3, 4; open nodes were not tested because an ancestor failed to reject the single-domain null.

Second, we tested whether SigDoTS retained a single domain when candidate domains were imposed within the interior of Layer 3 of section 151673 (Fig. 3h), a laminar compartment with limited evidence of discrete molecular substructure compared with other layers [14, 24, 25]. As a complementary assessment, we used cell2location with an external scRNA-seq reference to estimate spot-level cell-type proportions within the Layer 3 interior. The local and global composition-homogeneity scores were *H*_*l*_ = 0.966 and *H*_*g*_ = 0.977, respectively (Supplementary Methods S1.7; Fig. 3i). Values of *H*_*l*_ near 1 indicate that neighboring spots have similar cell-type compositions, whereas values of *H*_*g*_ near 1 indicate compositional similarity across the complete Layer 3 interior. We generated candidate domains with BANKSY at different levels of granularity *K*_0_> = 2, 3 and 4 (Fig. 3j). SigDoTS consistently retained one domain for *K*_0_> = 2, 3 and 4, with root *p*-values of 0.976, 0.976 and 1.000, respectively. Thus, the SigDoTS decision was invariant to candidate granularity, whereas HC-Wald retained every imposed split and scSHC retained two domains for *K*_0_> = 3 (Supplementary Results S2.2).

### SigDoTS separates epithelial and fibroblast domains that differ in immune-cell signals in ovarian cancer

To determine whether SigDoTS could retain molecularly distinct spatial domains beyond tissues with layered organization such as brain, we applied it to the SPATCH ovarian cancer FFPE Visium HD dataset [26] (Supplementary Methods S1.4). We selected a region of interest (ROI) containing multiple cell types. The ROI contained 13,690 8*μ*m bins (Fig. 4a,b). BANKSY generated *K*_0_> = 15 candidate domains in the ROI, equal to the number of cell types in the finest-resolution annotation from the original study (Fig. 4b). SigDoTS rejected the single-domain null at the root (*p* = 0.008) and merged them into six retained domains (Fig. 4c; Supplementary Fig. S4a,b). Using SPATCH transcriptomic labels transferred from matched single-cell RNA-sequencing data, we found that D1 and D2 were enriched for epithelial cells, D3 for macrophages and D4 for fibroblasts. D5 contained fibroblast, plasma-cell and T-cell signals, whereas D6 contained smooth-muscle and endothelial cells (Fig. 4d). To evaluate the RNA-defined domains in an orthogonal modality, we analyzed manually annotated single-cell CODEX measurements from an adjacent section [26]. After cross-section registration, we mapped each CODEX cell centroid to its nearest analyzed bin within 12 *μ*m and assigned each mapped cell the SigDoTS and BANKSY domain assignments of that bin. Pan-Cytokeratin was higher in D1–D2 and CD163 in D3 than in the remaining domains; SMA was higher in D4–D6 than in D1–D3, and CD34 was higher in D6 than in the remaining domains (Fig. 4f,g).

**Figure 4:**
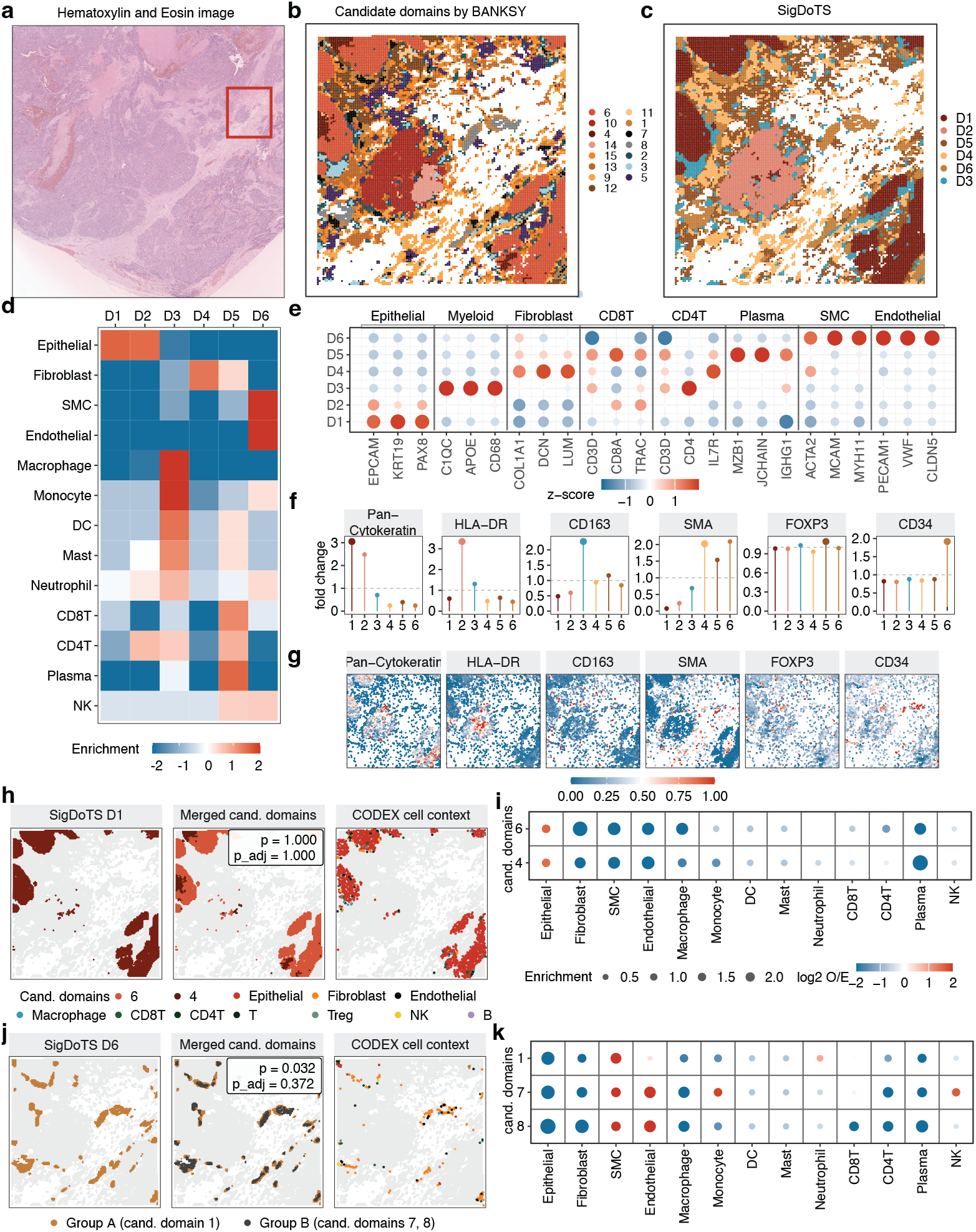
Six spatial domains retained by SigDoTS in ovarian cancer Visium HD data. **a** Hematoxylin-and-eosin-stained section; the box marks the analyzed region of interest (ROI). **b** Initial BANKSY candidate domains (*K*_0_> = 15) across 13,690 8 *μ*m bins. **c** Six domains retained by SigDoTS. **d** Enrichment of SPATCH-annotated cell types within each retained domain (log_2_ ratio of the within-domain to the ROI-wide proportion). **e** RNA-marker expression across retained domains (color, mean log-normalized expression standardized across domains for each gene; dot size, its absolute value). **f** Mean CODEX protein intensity in each domain relative to the other domains, measured in an adjacent tissue section; dashed lines mark a fold change of 1. **g** Spatial distributions of the CODEX proteins in **f. h**,**i** Retained domain D1, its candidate domains 4 and 6 with the corresponding CODEX cell-type map (**h**), and their SPATCH cell-type enrichment (**i**; color, log_2_ observed/expected enrichment; dot size, its absolute value). **j**,**k** Retained domain D6, the candidate split between domain 1 and domains 7 and 8 with the corresponding CODEX cell-type map (**j**), and SPATCH cell-type enrichment in candidate domains 1, 7 and 8 (**k**).

Specifically, SigDoTS separated two domains that were both epithelial-rich but differed in their immune-associated RNA and protein profiles. D1 had higher expression of the epithelial tumor-cell markers EPCAM, KRT19 and PAX8, whereas D2 had higher expression of the T-cell markers CD8A and TRAC and the highest HLA-DR protein signal among the six retained domains [27, 28] (Fig. 4e–g). A related distinction occurred between the fibroblast-containing domains: D4 was dominated by fibroblasts, whereas D5 also contained plasma-cell and T-cell signals.

To illustrate how SigDoTS merged the 15 BANKSY candidate domains into six retained domains, we examined two merges: D1, whose candidate split failed to reject the single-domain null, and D6, whose candidate split rejected it before but not after hierarchical adjustment. For D1, the unadjusted test failed to reject the single-domain null (*p* = 1.000), and candidate domains 4 and 6 were both enriched for epithelial cells (Fig. 4h,i); reporting them separately would have treated one epithelial region as two. For D6, the unadjusted test rejected the single-domain null (*p* = 0.032), so testing at *α* = 0.05 without adjustment would have retained its three candidate domains separately, but the hierarchically adjusted test (Methods) failed to reject it (*p*,_adj_ = 0.372; Supplementary Fig. S5a). These candidate domains shared a mural–endothelial profile but differed in composition: candidate domain 1 contained the largest smooth-muscle-cell fraction, whereas candidate domains 7 and 8 contained larger endothelial-cell fractions (Fig. 4j,k). Hierarchical adjustment thus kept this compositional variation as heterogeneity within one vascular domain rather than reporting it as three domains.

In a second ovarian cancer dataset profiled by standard Visium, SigDoTS retained nine domains from 15 BANKSY candidate domains and again separated a fibroblast-associated domain with higher T-cell and plasma-cell marker expression (D2) from fibroblast-associated domains with lower expression (D5–D7), while merging seven epithelial candidate domains that shared PAX8 and EPCAM expression (Supplementary Fig. S6; Supplementary Methods S1.5; Supplementary Results S2.2). The fibroblast–immune distinction therefore recurred across two platforms of different spatial resolution.

### SigDoTS distinguishes proliferative from mucin-associated epithelial domains in colorectal adenocarcinoma

We next applied SigDoTS to an ROI containing 25,343 cells from a colorectal adenocarcinoma Xenium 5K dataset [26] (Fig. 5a; Supplementary Methods S1.6). BANKSY generated *K*_0_> = 15 candidate domains. SigDoTS rejected the single-domain null at the root (*p* = 0.008) and merged them into seven retained domains (Fig. 5b; Supplementary Fig. S7a,b). Using SPATCH transcriptomic cell-type annotations, we found that D1, D4 and D5 were enriched for epithelial cells; D2 and D3 for plasma cells; D6 for CD4 and CD8 T cells, macrophages and neutrophils; and D7 for fibroblasts, endothelial cells and smooth-muscle cells (Fig. 5c). Adjacent-section CODEX measurements localized Pan-Cytokeratin primarily to D4 and D5, HLA-DR to D6 and SMA to D7 (Fig. 5e,f).

**Figure 5:**
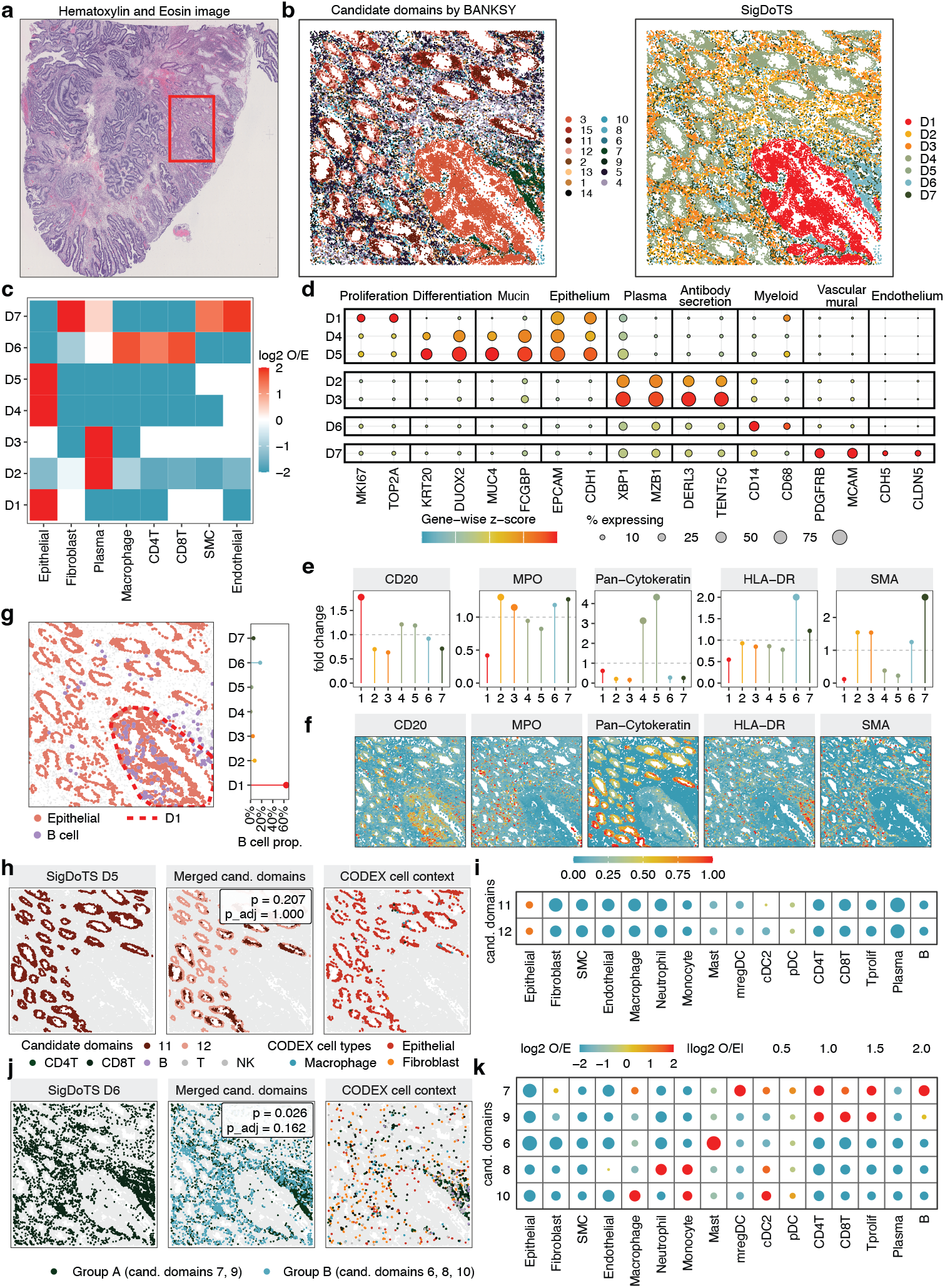
Seven spatial domains retained by SigDoTS in colorectal adenocarcinoma Xenium 5K data. **a** Hematoxylin-and-eosin-stained section; the box marks the analyzed region of interest (ROI). **b** Initial BANKSY candidate domains (*K*_0_> = 15; left) and the seven domains retained by SigDoTS (right). **c** Enrichment of SPATCH-annotated cell types within each retained domain (log_2_ ratio of the within-domain to the ROI-wide proportion). **d** RNA-marker expression across retained domains (color, gene-wise standardized mean expression; dot size, percentage of cells expressing each gene). **e** Mean CODEX protein intensity in each domain relative to the other domains, measured in an adjacent tissue section; dashed lines mark a fold change of 1. **f** Spatial distributions of the CODEX proteins in **e. g** CODEX epithelial and B cells relative to the boundary of D1 (left), and the percentage of all mapped B cells in each retained domain (right). **h**,**i** Retained domain D5, its candidate domains 11 and 12 with the corresponding CODEX cell-type map (**h**), and their SPATCH cell-type enrichment (**i**; color, log_2_ observed/expected enrichment; dot size, its absolute value). **j**,**k** Retained domain D6, the candidate split between domains 7, 9 and domains 6, 8, 10 with the corresponding CODEX cell-type map (**j**), and SPATCH cell-type enrichment in the five candidate domains (**k**).

Specifically, SigDoTS separated three epithelial-rich domains with different transcriptional programs and cellular contexts. D1 had higher proliferation-marker expression, whereas D4 and D5 had higher intestinal epithelial and mucin-associated marker expression, with the strongest mucin signal in D5 (Fig. 5d). Adjacent-section CODEX further showed that D1 contained the largest share of mapped B cells (Fig. 5g) [29].

Three merges illustrate how SigDoTS combined the 15 candidate domains into seven. For D5, the unadjusted test failed to reject the single-domain null (*p* = 0.207), and candidate domains 11 and 12 were both epithelial with similar SPATCH cell-type profiles and adjacent-section CODEX compositions (Fig. 5h,i); there was no evidence to report them as separate domains. For D6 and D7, the unadjusted tests rejected the single-domain null (*p* = 0.026 and 0.024), so testing without adjustment would have split both, but the hierarchically adjusted tests did not (*p*_adj_= 0.162 and 0.20; Supplementary Figs. S8and S9a; Supplementary Results S2.2). Within D6, candidate domains 7 and 9 were richer in CD4 and CD8 T cells, whereas candidate domains 6, 8 and 10 were enriched for mast cells, neutrophils and monocytes, and macrophages and monocytes, respectively (Fig. 5j,k); those within D7 differed in fibroblast content. SigDoTS thus kept this immune and fibroblast variation as heterogeneity within D6 and D7 rather than reporting it as separate domains.

## Methods

### The SigDoTS model

Let *s*_*i*_ ∈ ℝ ^2^ denote the two-dimensional coordinate of spatial unit *i*, and let 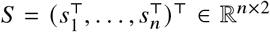 denote the spatial-coordinate matrix. We denote the corresponding preprocessed expression matrix used for spatial domain detection with *X* ∈ ℝ ^*n*×*p*^. SigDoTS models the gene expression matrix of a spatial domain as a matrix-normal distribution

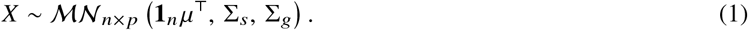

Here, *μ* ∈ ℝ^*p*^ is the mean vector of the *p* gene-expression features, assumed to be constant across spatial units under *H*_0_>. The matrix Σ_*s*_ ∈ ℝ^*n*×*n*^ is the spatial covariance matrix, whose entries depend on the spatial coordinates of the units, and Σ_*g*_ ∈ R^*p*×*p*^ is the gene–gene covariance matrix. Given two candidate spatial domains, SigDoTS tests whether the expression matrix within the merged region follows a single matrix-normal distribution (*H*_0_>: Eq. (1); *H*_1_: *X* does not follow any distribution of the above matrix-normal form). In this two-domain scenario, the merged region is the entire analyzed region; the multi-domain scenario below applies the same test to a node with *n*_*v*_ spatial units.

### Two-domain scenario hypothesis testing

Let *A* be a deterministic domain-detection approach that maps the expression matrix and spatial coordinates to a binary split 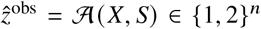, . To test the null hypothesis, SigDoTS uses the cluster index (CI) [15], the ratio of within-region gene-expression variation to total gene-expression variation; smaller values indicate greater separation. Given two candidate spatial domains *z* ∈ {1, 2}^*n*^, the CI is defined as

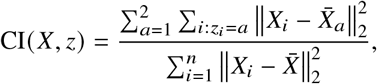

where 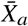 is the mean of cluster and 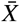 is the global mean. The two-domain SigDoTS procedure is therefore:

1. Run A on the observed data to obtain 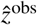 and compute 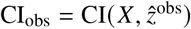.
2. Fit the spatial matrix-normal null in Eq. (1) to estimate 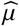, 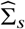and 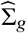 (see Null model estimation below).
3. Simulate null replicates 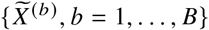 from the fitted null model (Supplementary Methods S1.1.4).
4. Rerun the identical pipeline 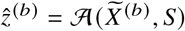, and compute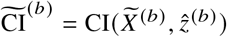.
5. Estimate the lower-tail Monte Carlo *p*-value 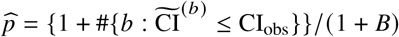

## Null model estimation

SigDoTS fits the null model in Eq. (1) to the tested region. Under the null hypothesis, SigDoTS estimates *μ* by the column means of *X*. The spatial covariance matrix Σ_*s*_ and the gene–gene covariance matrix Σ_*g*_ are nuisance parameters that describe variation within a domain. To prevent mean differences between true domains from appearing as spurious spatial and gene–gene covariance, which would reduce the power of the test, SigDoTS estimates these covariance matrices under a working model that allows a separate mean expression profile for each candidate domain,

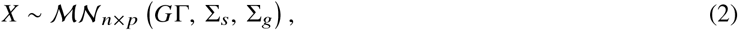

and plugs the estimates into the null model [30–32]. Here, *K*_*v*_ is the number of candidate domains in the tested region 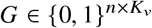, is the indicator matrix of candidate-domain membership, and row *k* of 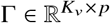 is the mean expression profile of candidate domain *k*. The working model uses the same covariance matrices as the null model and contains the null model as the special case in which all rows of Γare equal. When the tested region contains more than one true domain, the working model still holds provided that each candidate domain lies within one true domain. SigDoTS models the spatial covariance matrix Σ_*s*_ as Σ_*s*_ (*η,l*) = *ηKl*+ (1−*η*) *I*_*n*_, where *K*_*l*_ is an *n* × *n* Matérn -3/2 spatial kernel matrix, *I*_*n*_ is the identity matrix, *η* ∈[ 0, 1] is the spatial variance fraction, and *l* > 0 is the spatial length-scale parameter [33]. SigDoTS estimates and *l* under the working model, from pairs of spatial units that belong to the same candidate domain, by choosing the values for which the correlations predicted by the working model best match the observed correlations across distance classes (Supplementary Methods S1.1.3). SigDoTS then estimates Σ_*g*_ under the working model, with Σ_*s*_ fixed at 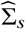: after removing the candidate-domain means and the fitted spatial dependence from the residuals, it assumes the spiked covariance model [15, 34–36] and estimates the number of detectable spikes and the residual variance iteratively (Supplementary Methods S1.1.3). SigDoTS factorizes 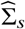 for whitening and null simulation by full eigendecomposition or with a Vecchia approximation [37, 38] (Supplementary Methods S1.1.5).

### Multi-domain scenario hypothesis testing

SigDoTS extends the two-domain test to multiple domains as follows (Supplementary Methods S1.1.6). For an internal node *v, n*_*v*_ denotes its number of spatial units. The multi-domain SigDoTS procedure is:

1. Build a binary tree over the *K*_0_> initial candidate domains by Ward-linkage clustering of spatially adjacent domains in principal-component space [17].
2. At each internal node *v*, apply the two-domain test described above, rerunning the *K*_*v*_-domain detection and tree construction on every null replicate and computing the null CI at the root split of each replicate tree.
3. Test from the root downward at 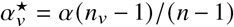, stopping below any node that fails to reject, which, when the nodewise *p*-values are valid, controls the family-wise error rate at level *α* [16, 39]. Equivalently, the hierarchically adjusted *p*-value 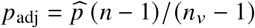 is compared with *α*.

## Discussion

SigDoTS is a post-clustering statistical framework for limiting oversegmentation in spatial domain detection. It complements existing domain-detection methods by testing which splits among their candidate domains to retain, under family-wise error control. Two features distinguish it from existing tests: rerunning the domain-detection method on each null replicate accounts for the selection of candidate domains from the tested data, and simulating the replicates from a matrix-normal single-domain null accounts for gene–gene covariance and spatial dependence.

Both features were necessary for type I error control. HC-Wald, which treats candidate domains as prespecified, rejected the global null in every dataset, and scSHC, which accounts for selection but assumes independent spatial units, rejected it in most datasets, increasingly so as spatial correlation strengthened. Under spatial dependence, clustering can divide smooth expression patterns that contain no mean differences, and null replicates simulated under independence do not reproduce this separation.

Two limitations follow from the same post-clustering design. It evaluates only boundaries among the initial candidate domains, so it cannot recover a biological boundary that the upstream method fails to propose, and it reruns that method on every null replicate, so its computational cost grows with the number of spatial units and with the runtime of that method.

## Code and Data Availability

The SigDoTS R package is available at https://github.com/Yan-Lab-MSU/SigDoTS; all datasets analyzed in this study are publicly available, with sources listed in Supplementary Data Availability S3.

