## Supplementary Material for "Statistical significance testing for spatial domain detection in spatial transcriptomics"

**S3 Data Availability** **17**

**S4 Supplementary Figures** **18**

### S1 Supplementary Methods

#### S1.1 The SigDoTS method

##### S1.1.1 The SigDoTS model and the two-domain test

We denote by $C \in \mathbb{N}_0^{n\times p}$ the gene-count matrix, where rows correspond to $n$ platform-specific spatial units (cells, bins, or multicellular spots) and columns correspond to $p$ genes, with $C_{ig}$ indicating the read or unique molecular identifier (UMI) count of gene $g$ in spatial unit $i$. Let $s_i \in \mathbb{R}^2$ denote the two-dimensional coordinate of spatial unit $i$, and let $S = (s_1^\top, \ldots, s_n^\top)^\top \in \mathbb{R}^{n\times 2}$ denote the spatial-coordinate matrix. We denote the corresponding preprocessed expression matrix used for spatial domain detection with $X \in \mathbb{R}^{n\times p}$. The matrix $X$ was obtained by library-size normalization followed by a $\log(1+x)$ transformation, with gene-wise centering and scaling applied when required by the upstream domain-detection method. Gene expression matrix $X$ will be used as input for the SigDoTS method.

SigDoTS models the gene expression matrix of a spatial domain as a matrix-normal distribution

$$X \sim \mathcal{MN}_{n\times p}\left(\mathbf{1}_n\mu^\top, \Sigma_s, \Sigma_g\right). \tag{S1}$$

Here, $\mu \in \mathbb{R}^p$ is the mean vector of the $p$ gene-expression features, assumed to be constant across spatial units under $H_0$. The matrix $\Sigma_s \in \mathbb{R}^{n\times n}$ is the spatial covariance matrix, whose entries depend on the spatial coordinates of the units, and $\Sigma_g \in \mathbb{R}^{p\times p}$ is the gene–gene covariance matrix. Therefore, given the two candidate spatial domains $\hat{z}^{\text{obs}}$, SigDoTS tests whether the expression matrix within the merged region follows a single matrix-normal distribution:

$$\begin{aligned} H_0: &\quad X \sim \mathcal{MN}_{n\times p}\left(\mathbf{1}_n\mu^\top, \Sigma_s, \Sigma_g\right), \\ &\quad \text{equivalently,} \quad \operatorname{vec}(X) \sim \mathcal{N}_{np}\left(\operatorname{vec}(\mathbf{1}_n\mu^\top), \Sigma_g \otimes \Sigma_s\right); \\ H_1: &\quad X \text{ does not follow any distribution of the above matrix-normal form.} \end{aligned}$$

The single-domain null assumes that all spatial measurement units within the tested region share a common mean gene-expression vector, while allowing expression to be correlated across genes and measurement units. The rationale is that neighboring measurement units may have correlated molecular profiles even when they belong to one spatial domain and share the same mean expression profile. By contrast, methods for testing cluster significance commonly model observations as independent and identically distributed multivariate random vectors. For example, scSHC assumes $X_{i\cdot} \overset{\text{i.i.d.}}{\sim} \mathcal{N}_p(\mu, \Sigma_g),\ i = 1, \ldots, n$, which corresponds to the special case $\Sigma_s = I_n$ under the matrix-normal formulation. SigDoTS instead treats the expression matrix within the tested region as one realization from a spatially correlated matrix-normal model. The covariance matrix $\Sigma_g$ represents dependence among genes, whereas $\Sigma_s$ represents dependence among spatial measurement units as a function of their locations. This formulation tests differences in mean expression between candidate domains while including within-domain spatial covariance in the null model. When $H_0$ is not rejected, the data do not provide sufficient evidence against the single-domain matrix-normal model, and SigDoTS does not retain the candidate split.

**Two-domain scenario hypothesis testing** We start by introducing the two-domain scenario where we test whether a spatial region should be split into two spatial domains. Let $\mathcal{A}$ be a deterministic domain-detection approach that maps the expression matrix and spatial coordinates to a binary split, $\hat{z}^{\text{obs}} = \mathcal{A}(X, S) \in \{1, 2\}^n$. To test the null hypothesis, SigDoTS uses the cluster index (CI) [1], the ratio of within-region gene-expression variation to total gene-expression variation; smaller values indicate greater separation. Given two candidate spatial domains $z \in \{1, 2\}^n$, the CI is defined as

$$\mathrm{CI}(X, z) = \frac{\sum_{a=1}^{2} \sum_{i:z_i=a} \left\|X_i - \bar{X}_a\right\|_2^2}{\sum_{i=1}^{n} \left\|X_i - \bar{X}\right\|_2^2},$$

where $\bar{X}_a$ is the mean of cluster $a$ and $\bar{X}$ is the global mean.

Given the pre-processed gene expression matrix $X$, spatial coordinates $S$ and a deterministic domain detection pipeline $\mathcal{A}$, the two-domain SigDoTS procedure is therefore:

1. Run $\mathcal{A}$ on the observed data to obtain $\hat{z}^{\text{obs}}$ and compute $\mathrm{CI}_{\text{obs}} = \mathrm{CI}(X, \hat{z}^{\text{obs}})$.

2. Fit the spatial matrix-normal null in Eq. (S1) to estimate $\widehat{\mu}$, $\widehat{\Sigma}_s$ and $\widehat{\Sigma}_g$.

3. Simulate null replicates $\{\widetilde{X}^{(b)}, b = 1, \dots, B\}$ from the fitted null model.
4. Rerun the identical pipeline, $\hat{z}^{(b)} = \mathcal{A}(\widetilde{X}^{(b)}, S)$, and compute $\widetilde{\mathrm{CI}}^{(b)} = \mathrm{CI}(\widetilde{X}^{(b)}, \hat{z}^{(b)})$.
5. Estimate the lower-tail Monte Carlo $p$-value

$$\widehat{p} = \frac{1 + \#\{b : \widetilde{\mathrm{CI}}^{(b)} \leq \mathrm{CI}_{\mathrm{obs}}\}}{1 + B}.$$

A small $p$-value indicates that the binary split produced by the pipeline is more separated than expected under a single spatially coherent null with the fitted covariance matrices.

##### S1.1.2 Matrix-normal properties

**Definition 1** (Matrix-normal distribution). *A random matrix $X \in \mathbb{R}^{n\times p}$ follows a matrix-normal distribution*

$$X \sim \mathcal{MN}_{n\times p}(M, \Sigma_s, \Sigma_g)$$

*if and only if*

$$\mathrm{vec}(X) \sim \mathcal{N}\big(\mathrm{vec}(M),\ \Sigma_g \otimes \Sigma_s\big),$$

*where $\Sigma_s \in \mathbb{R}^{n\times n}$ is the row (spatial) covariance and $\Sigma_g \in \mathbb{R}^{p\times p}$ is the column (gene) covariance.*

A direct consequence of the Kronecker covariance is that entrywise covariances factorize:

$$\mathrm{Cov}(X_{ia}, X_{jb}) \ = \ (\Sigma_s)_{ij}\,(\Sigma_g)_{ab}.$$

Equivalently, the cross-covariance between entire rows is

$$\mathrm{Cov}\big(X_{i\cdot}, X_{j\cdot}\big) \ = \ (\Sigma_s)_{ij}\,\Sigma_g, \tag{S2}$$

and similarly the cross-covariance between entire columns is

$$\mathrm{Cov}\big(X_{\cdot a}, X_{\cdot b}\big) \ = \ (\Sigma_g)_{ab}\,\Sigma_s.$$

Thus, under separability, **all row–row dependence is carried by** $\Sigma_s$ (as a scalar multiplier), and **all column–column dependence is carried by** $\Sigma_g$ (as a shared covariance "shape").

**Lemma 1** (Affine transformation of a matrix-normal random matrix). *If $X \sim \mathcal{MN}(M, \Sigma_s, \Sigma_g)$ and $A \in \mathbb{R}^{n'\times n}$, $B \in \mathbb{R}^{p'\times p}$ are fixed matrices, then*

$$AXB^\top \sim \mathcal{MN}\big(AMB^\top,\ A\Sigma_s A^\top,\ B\Sigma_g B^\top\big),$$

*whenever the resulting covariances are well-defined.*

Assume the null is correctly specified:

$$X \sim \mathcal{MN}\big(\mathbf{1}_n\mu^\top,\ \Sigma_s,\ \Sigma_g\big).$$

Let $X_c = X - \mathbf{1}_n\mu^\top$, so

$$X_c \sim \mathcal{MN}(0, \Sigma_s, \Sigma_g).$$

Define the row-whitened matrix

$$Y := \Sigma_s^{-1/2} X_c.$$

**Proposition 1** (Whitening makes rows i.i.d.). *If $X_c \sim \mathcal{MN}(0, \Sigma_s, \Sigma_g)$ and $\Sigma_s \succ 0$, then*

$$Y = \Sigma_s^{-1/2} X_c \sim \mathcal{MN}(0, I_n, \Sigma_g).$$

*In particular, the rows of $Y$ are i.i.d.:*

$$Y_{i,\cdot} \overset{i.i.d.}{\sim} \mathcal{N}_p(0, \Sigma_g).$$

*Proof.* Apply the linear transformation lemma with $A = \Sigma_s^{-1/2}$ and $B = I_p$:

$$Y = \Sigma_s^{-1/2} X_c \sim \mathcal{MN}\big(0,\ \Sigma_s^{-1/2}\Sigma_s\Sigma_s^{-1/2},\ \Sigma_g\big) = \mathcal{MN}(0, I_n, \Sigma_g).$$

The i.i.d. row statement follows from the definition of $\mathcal{MN}(0, I_n, \Sigma_g)$. □

##### S1.1.3 Null model estimation

SigDoTS fits the null model in Eq. (S1) to the tested region. The mean vector $\mu$ is estimated under the null hypothesis. The spatial covariance matrix $\Sigma_s$ and the gene–gene covariance matrix $\Sigma_g$ are nuisance parameters: they describe the variation within a domain and are not the subject of the test. SigDoTS estimates them under a working model that allows a separate mean expression profile for each candidate domain,

$$X \sim \mathcal{MN}_{n\times p}\left(G\Gamma,\ \Sigma_s(\eta,\ell),\ \Sigma_g\right), \tag{S3}$$

and plugs the estimates into the null model [2–4]. Here, $K_v$ is the number of candidate domains in the tested region, and $G \in \{0,1\}^{n\times K_v}$ records the candidate spatial domain assignment of each spatial unit, with $G_{ik} = 1$ if spatial unit $i$ belongs to candidate domain $k$ and $G_{ik} = 0$ otherwise, and row $k$ of $\Gamma \in \mathbb{R}^{K_v\times p}$ is the mean expression profile of candidate domain $k$. For the two-domain test, the working model contains the two observed candidate domains. The working model uses the same covariance matrices as the null model and contains the null model as the special case in which all rows of $\Gamma$ are equal. When the null hypothesis holds, the plug-in estimates therefore estimate the covariance matrices of the null model. When the tested region contains more than one true domain, the working model still holds provided that each candidate domain lies within one true domain. Between-domain mean differences are then captured by ($\Gamma$) rather than by the covariance estimates. Estimating the covariance matrices under the null model would instead attribute these mean differences to within-domain variation, introducing spurious spatial and gene–gene covariance: the test would lose power. The null replicates are drawn from the fitted null model $\mathcal{MN}_{n\times p}(\mathbf{1}_n\widehat{\mu}^\top, \widehat{\Sigma}_s, \widehat{\Sigma}_g)$. SigDoTS obtains $\widehat{\mu}$, $\widehat{\Sigma}_s$ and $\widehat{\Sigma}_g$ in three steps.

**Estimation of mean vector $\mu$** Under the null hypothesis, all spatial units share the mean vector $\mu$. SigDoTS estimates $\mu$ by the column means of $X$, $\widehat{\mu} = n^{-1}X^\top\mathbf{1}_n$, and denotes the centered expression matrix by $X_c = X - \mathbf{1}_n\widehat{\mu}^\top$.

**Estimation of spatial covariance matrix $\Sigma_s$** SigDoTS models the spatial covariance matrix $\Sigma_s$ as

$$\Sigma_s(\eta,\ell) = \eta K_\ell + (1-\eta)I_n, \tag{S4}$$

where $K_\ell$ is an $n\times n$ spatial kernel matrix, $I_n$ is the identity matrix, $\eta \in [0,1]$ is the spatial variance fraction, and $\ell > 0$ is the spatial length-scale parameter. Specifically, SigDoTS uses a Matérn kernel with smoothness parameter $\nu = 3/2$,

$$K_\ell(s_i,s_j) = \left(1 + \frac{\sqrt{3}\,\|s_i - s_j\|_2}{\ell}\right)\exp\left(-\frac{\sqrt{3}\,\|s_i - s_j\|_2}{\ell}\right).$$

The kernel satisfies $K_\ell(s_i,s_i) = 1$, giving $\Sigma_s(\eta,\ell)$ unit diagonal. Thus, $\eta$ is the fraction of marginal variance assigned to the spatially correlated component, and $1-\eta$ is the fraction assigned to the spatially uncorrelated component. For distinct spatial units $i$ and $j$, the correlation is $\eta K_\ell(s_i,s_j)$. At fixed $\ell$, $\eta$ scales these correlations; for $\eta > 0$, a larger $\ell$ makes them decay more slowly with distance.

SigDoTS estimates $\eta$ and $\ell$ under the working model, from pairs of spatial units that belong to the same candidate domain. Before the estimation, SigDoTS subtracts the mean of each gene within each candidate domain, which removes the term $G\Gamma$ of the working model. SigDoTS sorts the pairs into distance classes, so that the two units of every pair in a class are about equally far apart. For distance class $c$, $N_c$ denotes the number of pairs, and $\widehat{\rho}_c$ denotes the correlation between the expression values of the two units in a pair, computed from all genes together. For given $\eta$ and $\ell$, $\rho_c(\eta,\ell)$ denotes the value of $\widehat{\rho}_c$ predicted by the working model. This predicted value depends on neither $\Gamma$ nor $\Sigma_g$. SigDoTS chooses the values of $\eta$ and $\ell$ for which the predicted correlations best match the observed correlations, with more weight on distance classes that contain more pairs:

$$(\widehat{\eta},\widehat{\ell}) = \operatorname*{argmin}_{\eta,\ell}\sum_c N_c\left\{\widehat{\rho}_c - \rho_c(\eta,\ell)\right\}^2.$$

The paragraph "Spatial parameters" below gives the formulas for $\widehat{\rho}_c$ and $\rho_c(\eta,\ell)$. The fitted spatial covariance matrix is $\widehat{\Sigma}_s = \Sigma_s(\widehat{\eta},\widehat{\ell}) + \tau I_n$, where the small constant $\tau = 10^{-6}$ keeps $\widehat{\Sigma}_s$ numerically invertible.

**Estimation of gene-gene covariance matrix $\Sigma_g$** SigDoTS then estimates $\Sigma_g$ under the working model, with $\Sigma_s$ fixed at $\widehat{\Sigma}_s$. As for $\eta$ and $\ell$, SigDoTS first subtracts the mean of each gene within each candidate domain. Because $\widehat{\Sigma}_s$ is now available, these means are estimated by generalized least squares, which accounts for the correlation between spatial units. SigDoTS then removes the fitted spatial dependence from the residuals and writes the result as a matrix $Y$ with $n - K_v$ rows, one fewer for each of the $K_v$ candidate-domain means removed. Under the working model, the rows of $Y$ are approximately independent, and each row has covariance matrix $\Sigma_g$. Because $Y$ has $n - K_v$ rows, $n$ is replaced by $n - K_v$ in the four steps below.

The number of genes can be comparable to, or larger than, the number of rows of $Y$, so the ordinary sample covariance matrix $n^{-1}Y^\top Y$ is an unreliable estimate of $\Sigma_g$. SigDoTS therefore estimates $\Sigma_g$ from the eigenvalues and eigenvectors of $n^{-1}Y^\top Y$ in four steps. First, SigDoTS assumes the spiked covariance model [1, 5–7]

$$\Sigma_g = V_r \Theta_r V_r^\top + \sigma^2 \left(I_p - V_r V_r^\top\right), \tag{S5}$$

where $r$ is the number of population spikes, the columns of $V_r$ are the corresponding orthonormal eigenvectors, $\Theta_r = \operatorname{diag}(\theta_1, \dots, \theta_r)$ contains the spike eigenvalues with $\theta_1 \geq \cdots \geq \theta_r > \sigma^2$, and $\sigma^2$ is the common eigenvalue in the remaining $p - r$ directions. SigDoTS estimates this model by

$$\widehat{\Sigma}_g = \widehat{V}_{\widehat{r}} \widehat{\Theta}_{\widehat{r}} \widehat{V}_{\widehat{r}}^\top + \widehat{\sigma}^2 \left(I_p - \widehat{V}_{\widehat{r}} \widehat{V}_{\widehat{r}}^\top\right), \tag{S6}$$

where $\widehat{r}$ is the estimated number of detectable spikes, the columns of $\widehat{V}_{\widehat{r}}$ are the corresponding gene-loading directions, $\widehat{\Theta}_{\widehat{r}}$ contains the estimated spike eigenvalues, and $\widehat{\sigma}^2$ is the variance in the remaining $p - \widehat{r}$ directions.

Second, let $q = \operatorname{rank}(Y)$, and let $s_1 \geq \cdots \geq s_q > 0$ denote the nonzero singular values of $Y$. The corresponding nonzero eigenvalues of the sample covariance matrix $n^{-1}Y^\top Y$ are

$$\widehat{\lambda}_j = \frac{s_j^2}{n}, \qquad j = 1, \dots, q. \tag{S7}$$

The right singular vector associated with $\widehat{\lambda}_j$ defines the gene-loading direction for the $j$th sample eigenvalue.

Third, SigDoTS estimates the number of detectable spikes and the residual variance iteratively. Given a current estimate of $\sigma^2$, SigDoTS first updates the number of detectable spikes. With $\gamma = p/n$, the sample covariance eigenvalues corresponding to the common population eigenvalue $\sigma^2$ are asymptotically bounded above by $\lambda_+(\sigma^2) = \sigma^2(1 + \sqrt{\gamma})^2$, whereas sufficiently large population spikes produce sample eigenvalues above this bound [5]; SigDoTS therefore defines a spike by the criterion $\widehat{\lambda}_j \geq \lambda_+(\widehat{\sigma}^2)$ and estimates the number of spikes as

$$\widehat{r} = \min\left\{r_{\max}, \#\left\{j : \widehat{\lambda}_j \geq \lambda_+(\widehat{\sigma}^2)\right\}\right\}. \tag{S8}$$

Here, $r_{\max}$ is a prespecified upper bound. Given the updated value of $\widehat{r}$, SigDoTS updates the residual variance $\widehat{\sigma}^2$. For a candidate value $\sigma^2$, we apply the Wang–Fan bias correction [8] by defining the corrected estimate of the $j$th spike eigenvalue as

$$\widetilde{\theta}_j(\sigma^2) = \widehat{\lambda}_j - \sigma^2 \frac{p}{n}, \qquad j = 1, \dots, \widehat{r}, \tag{S9}$$

SigDoTS chooses $\widehat{\sigma}^2$ so that the $\widehat{r}$ corrected spike eigenvalues and the $p - \widehat{r}$ residual eigenvalues sum to the total sample variance:

$$\sum_{j=1}^{q} \widehat{\lambda}_j = \sum_{j=1}^{\widehat{r}} \widetilde{\theta}_j(\widehat{\sigma}^2) + (p - \widehat{r})\widehat{\sigma}^2. \tag{S10}$$

Substituting equation (S9) into equation (S10) and solving for $\widehat{\sigma}^2$ gives

$$\widehat{\sigma}^2 = \frac{\sum_{j=1}^{q} \widehat{\lambda}_j - \sum_{j=1}^{\widehat{r}} \widehat{\lambda}_j}{p - \widehat{r} - p\widehat{r}/n}. \tag{S11}$$

Then SigDoTS alternates between the spike-count and residual-variance updates until convergence.

Finally, after convergence, SigDoTS estimates each spike eigenvalue as

$$\widehat{\theta}_j = \max\left\{\widehat{\lambda}_j - \widehat{\sigma}^2 \frac{p}{n},\ 0\right\}, \qquad j = 1, \ldots, \widehat{r}. \tag{S12}$$

The maximum with zero ensures that the estimated covariance eigenvalues are nonnegative. SigDoTS then sets $\widehat{\Theta}_{\widehat{r}} = \operatorname{diag}(\widehat{\theta}_1, \ldots, \widehat{\theta}_{\widehat{r}})$, and the columns of $\widehat{V}_{\widehat{r}}$ are the corresponding right singular vectors of $Y$. If $\widehat{r} = 0$, the estimator reduces to $\widehat{\Sigma}_g = \widehat{\sigma}^2 I_p$.

**Spatial parameters** Let $K_v$ be the number of candidate domains in the tested region and let $P_G = I_n - G(G^\top G)^{-1}G^\top$, with $G$ the $n \times K_v$ candidate-domain indicator matrix. The matrix $\widetilde{X} = P_G X_c$ contains the expression values after the mean of each gene within each candidate domain has been subtracted. Let $\mathcal{P}_c$ denote the pairs of distinct spatial units in the same candidate domain that fall in distance class $c$, and let $N_c = |\mathcal{P}_c|$. The observed and predicted correlations in distance class $c$ are

$$\widehat{\rho}_c = \frac{N_c^{-1} \sum_{(i,i') \in \mathcal{P}_c} (\widetilde{X}\widetilde{X}^\top)_{ii'}}{n^{-1} \operatorname{tr}(\widetilde{X}\widetilde{X}^\top)}, \qquad \rho_c(\eta, \ell) = \frac{N_c^{-1} \sum_{(i,i') \in \mathcal{P}_c} \{P_G \Sigma_s(\eta, \ell) P_G\}_{ii'}}{n^{-1} \operatorname{tr}\{P_G \Sigma_s(\eta, \ell) P_G\}}.$$

Here, $(\widetilde{X}\widetilde{X}^\top)_{ii'} = \sum_{j=1}^{p} \widetilde{X}_{ij}\widetilde{X}_{i'j}$ sums over all genes. Under the working model in Eq. (S3) with fixed candidate domains, $\mathbb{E}(\widetilde{X}\widetilde{X}^\top) = \operatorname{tr}(\Sigma_g)\, P_G \Sigma_s P_G$ for every $\Gamma$ (section "Matrix-normal properties"). Thus, $\rho_c(\eta, \ell)$ is the ratio of the expected numerator and the expected denominator of $\widehat{\rho}_c$. The factor $\operatorname{tr}(\Sigma_g)$ cancels, so $\rho_c(\eta, \ell)$ depends on neither $\Sigma_g$ nor $\Gamma$.

**Removing the spatial dependence and the candidate-domain means** Under the working model in Eq. (S3), with $\Sigma_s$ fixed at $\widehat{\Sigma}_s$,

$$X = G\Gamma + E, \qquad E \sim \mathcal{MN}_{n\times p}(0, \widehat{\Sigma}_s, \Sigma_g).$$

Each column of $X$, the expression values of one gene, follows a linear regression on the $K_v$ columns of $G$ whose errors have covariance proportional to $\widehat{\Sigma}_s$. Let $\widehat{\Sigma}_s = \widehat{U}_s \widehat{\Lambda}_s \widehat{U}_s^\top$ be the eigendecomposition of $\widehat{\Sigma}_s$, with eigenvalues below a small numerical tolerance set to that tolerance, and let $\widehat{\Sigma}_s^{-1/2} = \widehat{U}_s \widehat{\Lambda}_s^{-1/2} \widehat{U}_s^\top$. Multiplying the model by $\widehat{\Sigma}_s^{-1/2}$ gives

$$X_w = G_w \Gamma + E_w, \qquad X_w = \widehat{\Sigma}_s^{-1/2} X, \quad G_w = \widehat{\Sigma}_s^{-1/2} G, \quad E_w = \widehat{\Sigma}_s^{-1/2} E \sim \mathcal{MN}_{n\times p}(0, I_n, \Sigma_g),$$

in which the rows of the errors are independent. The generalized least-squares estimator of the candidate-domain means $\Gamma$ is the ordinary least-squares estimator in this whitened model,

$$\widehat{\Gamma} = (G_w^\top G_w)^{-1} G_w^\top X_w = (G^\top \widehat{\Sigma}_s^{-1} G)^{-1} G^\top \widehat{\Sigma}_s^{-1} X,$$

and is the same for every gene. The whitened residuals are

$$X_w - G_w \widehat{\Gamma} = (I_n - H_w) X_w, \qquad H_w = G_w (G_w^\top G_w)^{-1} G_w^\top.$$

The matrix $I_n - H_w$ projects onto the orthogonal complement of the column space of $G_w$, which has dimension $n - K_v$. Let the columns of $Q \in \mathbb{R}^{n\times(n-K_v)}$ form an orthonormal basis of this complement. Then $I_n - H_w = QQ^\top$, and the whitened residuals equal $QY$, with

$$Y = Q^\top X_w = Q^\top \widehat{\Sigma}_s^{-1/2} X.$$

The $n$ rows of the whitened residuals are correlated, because their row covariance is $I_n - H_w$. In contrast, if the working model holds with known $\Sigma_s$ and fixed candidate domains, then $Y \sim \mathcal{MN}_{(n-K_v)\times p}(0, I_{n-K_v}, \Sigma_g)$ (section "Matrix-normal properties"), so the $n - K_v$ rows of $Y$ are independent $\mathcal{N}_p(0, \Sigma_g)$ random vectors. Both carry the same sums of squares and cross-products,

$$Y^\top Y = X^\top \left\{ \widehat{\Sigma}_s^{-1} - \widehat{\Sigma}_s^{-1} G \left( G^\top \widehat{\Sigma}_s^{-1} G \right)^{-1} G^\top \widehat{\Sigma}_s^{-1} \right\} X,$$

which does not depend on the choice of $Q$, and $\mathbb{E}(Y^\top Y) = (n - K_v)\Sigma_g$. Because $Q^\top G_w = 0$ and $\mathbf{1}_n = G\mathbf{1}_{K_v}$, $Y$ is unchanged when $X$ is replaced by $X_c$. In practice, SigDoTS computes the full QR decomposition $G_w = [\, Q_1 \;\; Q \,] \begin{pmatrix} R \\ 0 \end{pmatrix}$, where $[\,Q_1 \;\; Q\,]$ is an $n \times n$ orthogonal matrix and $R \in \mathbb{R}^{K_v \times K_v}$ is upper triangular, as in standard least-squares regression. The columns of $Q_1$ span the column space of $G_w$, so $H_w = Q_1 Q_1^\top$. SigDoTS applies $Q^\top$ through the Householder reflections of this decomposition without forming $Q$. With the Vecchia approximation, $\widehat{\Sigma}_s^{-1/2}$ is replaced by the approximate inverse Cholesky factor (section "Vecchia approximation for scalable spatial covariance operations"), and the same derivation applies.

**Computation** SigDoTS computes $\widehat{\Sigma}_s^{-1/2}$, and the factor of $\widehat{\Sigma}_s$ used to simulate the null replicates, either from the eigendecomposition of the full $n \times n$ matrix $\widehat{\Sigma}_s$ or with a Vecchia approximation [9, 10]. The eigendecomposition requires the evaluation and storage of all pairwise spatial covariances, resulting in $O(n^2)$ memory and $O(n^3)$ time. The Vecchia approximation, described in Section S1.1.5, performs the same transformations without constructing or eigendecomposing the full covariance matrix and is the default in the R package. When the full-matrix computation is requested, SigDoTS stops with an error if its estimated peak memory exceeds a user-specified limit $M_{\max}$ (16 GiB by default).

##### S1.1.4 Null data sampling

SigDoTS generates Monte Carlo replicates from the fitted null model

$$\widetilde{X} \sim \mathcal{MN}_{n\times p}\left(\mathbf{1}_n \widehat{\mu}^\top, \widehat{\Sigma}_s, \widehat{\Sigma}_g\right). \tag{S13}$$

Specifically, let $\widehat{\Sigma}_s = \widehat{U}_s \widehat{\Lambda}_s \widehat{U}_s^\top$ and $\widehat{\Sigma}_g = \widehat{V}_r \widehat{D}_r \widehat{V}_r^\top + \widehat{\sigma}^2 I_p$, SigDoTS draws independent matrices $W_0 \in \mathbb{R}^{n\times p}$ and $W_r \in \mathbb{R}^{n\times r}$ with i.i.d. $\mathcal{N}(0,1)$ entries and generates

$$\widetilde{X} = \mathbf{1}_n \widehat{\mu}^\top + \widehat{U}_s \widehat{\Lambda}_s^{1/2} W_0 \widehat{\sigma} + \widehat{U}_s \widehat{\Lambda}_s^{1/2} W_r \widehat{D}_r^{1/2} \widehat{V}_r^\top.$$

This formulation generates the fitted matrix-normal distribution in Equation (S13); the derivation follows.

**Derivation.** Let

$$\widehat{A}_s = \widehat{U}_s \widehat{\Lambda}_s^{1/2}, \qquad \widehat{\Sigma}_s = \widehat{A}_s \widehat{A}_s^\top,$$

and write

$$\widetilde{X} = \mathbf{1}_n \widehat{\mu}^\top + Z_0 + Z_r,$$

where

$$Z_0 = \widehat{A}_s W_0 \widehat{\sigma}, \qquad Z_r = \widehat{A}_s W_r \widehat{D}_r^{1/2} \widehat{V}_r^\top.$$

Because $W_0$ and $W_r$ have independent standard-normal entries,

$$\mathrm{vec}(W_0) \sim \mathcal{N}_{np}(0, I_{np}), \qquad \mathrm{vec}(W_r) \sim \mathcal{N}_{nr}(0, I_{nr}),$$

and the two random vectors are independent.

Using the identity

$$\mathrm{vec}(ABC) = (C^\top \otimes A)\,\mathrm{vec}(B),$$

we obtain

$$\mathrm{vec}(Z_0) = \widehat{\sigma}\left(I_p \otimes \widehat{A}_s\right)\mathrm{vec}(W_0).$$

Therefore,

$$\begin{aligned}\mathrm{Cov}\{\mathrm{vec}(Z_0)\} &= \widehat{\sigma}^2 \left(I_p \otimes \widehat{A}_s\right)\left(I_p \otimes \widehat{A}_s\right)^\top \\ &= \widehat{\sigma}^2 I_p \otimes \widehat{\Sigma}_s.\end{aligned} \tag{S14}$$

Similarly,

$$\mathrm{vec}(Z_r) = \left(\widehat{V}_r \widehat{D}_r^{1/2} \otimes \widehat{A}_s\right)\mathrm{vec}(W_r),$$

and hence

$$\begin{aligned}\mathrm{Cov}\{\mathrm{vec}(Z_r)\} &= \left(\widehat{V}_r\widehat{D}_r^{1/2}\otimes\widehat{A}_s\right)\left(\widehat{V}_r\widehat{D}_r^{1/2}\otimes\widehat{A}_s\right)^\top\\ &= \left(\widehat{V}_r\widehat{D}_r\widehat{V}_r^\top\right)\otimes\widehat{\Sigma}_s. \end{aligned}\tag{S15}$$

Because $W_0$ and $W_r$ are independent, the cross-covariance between $\mathrm{vec}(Z_0)$ and $\mathrm{vec}(Z_r)$ is zero. Consequently,

$$\begin{aligned}\mathrm{Cov}\{\mathrm{vec}(\widetilde{X})\} &= \mathrm{Cov}\{\mathrm{vec}(Z_0)\} + \mathrm{Cov}\{\mathrm{vec}(Z_r)\}\\ &= \left(\widehat{\sigma}^2 I_p + \widehat{V}_r\widehat{D}_r\widehat{V}_r^\top\right)\otimes\widehat{\Sigma}_s\\ &= \widehat{\Sigma}_g\otimes\widehat{\Sigma}_s.\end{aligned}\tag{S16}$$

Moreover,

$$\mathbb{E}(\widetilde{X}) = \mathbf{1}_n\widehat{\mu}^\top.$$

Since $\widetilde{X}$ is an affine transformation of jointly Gaussian random matrices, $\mathrm{vec}(\widetilde{X})$ is multivariate normal. Therefore,

$$\mathrm{vec}(\widetilde{X}) \sim \mathcal{N}_{np}\left(\mathrm{vec}(\mathbf{1}_n\widehat{\mu}^\top),\ \widehat{\Sigma}_g\otimes\widehat{\Sigma}_s\right),$$

which is equivalent to

$$\widetilde{X} \sim \mathcal{MN}_{n\times p}\left(\mathbf{1}_n\widehat{\mu}^\top,\ \widehat{\Sigma}_s,\ \widehat{\Sigma}_g\right).$$

##### S1.1.5 Vecchia approximation for scalable spatial covariance operations

Exact spatial whitening and simulation require the full $n\times n$ matrix $\widehat{\Sigma}_s$ and its dense eigendecomposition. This representation gives direct matrix square-root and inverse-square-root operations, but its memory and time costs grow quadratically or cubically with the number of spatial units. In large tissue regions, this cost can dominate the null-model estimation and Monte Carlo simulation. SigDoTS therefore uses the exact dense representation when the estimated memory requirement remains within a prespecified limit and uses a Vecchia approximation when the dense representation would exceed this limit, when dense factorization fails numerically, or when the analyst explicitly requests the approximation. In the current implementation, the Vecchia representation applies to the Matérn-3/2 spatial kernel.

The Vecchia approximation replaces the joint Gaussian density over all spatial units with a product of low-dimensional conditional Gaussian densities [9–11]. Let $z = (z_1,\dots,z_n)^\top$ denote one centered feature vector under the spatial null, after ordering the spatial locations. The exact density factorizes as $p(z)=\prod_{i=1}^n p(z_i\mid z_1,\dots,z_{i-1})$. Vecchia's approximation restricts each conditioning set to a small ordered neighbor set $N_i\subset\{1,\dots,i-1\}$:

$$p(z)\approx\prod_{i=1}^n p\left(z_i\,\middle|\,z_{N_i}\right).\tag{S17}$$

SigDoTS orders spatial units by a max–min rule, which spreads early units across the tissue region, and then assigns each unit its nearest previously ordered neighbors. This construction preserves local spatial dependence while keeping each conditional covariance calculation low-dimensional.

The conditional factors define a sparse inverse-Cholesky operator. SigDoTS uses this operator for the two operations required by the null model. For spatial whitening, it applies the inverse-Cholesky operator to each centered feature vector, which approximates multiplication by $\widehat{\Sigma}_s^{-1/2}$. For null simulation, it applies the corresponding spatial correlation operator to independent Gaussian noise, which approximates multiplication by $\widehat{\Sigma}_s^{1/2}$. Thus, the Vecchia approximation does not change the fitted single-domain null model; it changes how SigDoTS carries out the row-covariance operations when dense linear algebra becomes computationally impractical.

##### S1.1.6 Multi-domain scenario hypothesis testing

SigDoTS extends the two-domain test to multiple domains by organizing the initial candidate domains into a binary hierarchy. It tests the split at each visited internal node against a node-specific single-mean null and prunes the hierarchy through top-down testing with hierarchical adjustment. The procedure comprises three stages: construction of a spatially constrained domain tree, nodewise testing, and hierarchical error control.

**Stage 1: Construction of a spatially constrained domain tree** SigDoTS first applies a fixed, deterministic domain-detection pipeline $\mathcal{A}_{K_0}$ to the observed gene expression matrix $X$ and spatial coordinates $S$, producing $K_0 \geq 2$ candidate spatial domains. The $K_0$ candidate domains form the leaves of the observed hierarchy.

SigDoTS restricts tree construction to domains that are adjacent in space. It constructs a $k$-nearest-neighbor graph from the spatial coordinates, with default $k = 6$, and identifies the connected components within each leaf domain. A component enters the adjacency calculation when it contains at least an $\phi_{\min}$ fraction of the spatial units assigned to that domain. Components below this threshold remain assigned to their domains, but they cannot establish merge eligibility. Two leaf domains are adjacent when a retained component from one shares a graph edge with a retained component from the other. Two groups of leaf domains are adjacent when they contain at least one adjacent pair of leaf domains.

Following scSHC [12], SigDoTS applies hierarchical clustering with Ward linkage to the first 30 principal components of $X$ by default. In contrast to scSHC, SigDoTS begins with detected spatial domains rather than individual observations and restricts merges to adjacent domain groups. Let $Z^{\mathrm{PCA}}$ denote the retained principal-component scores. For a group $A$, let $n_A$ denote its number of spatial units and let $\overline{Z}_A^{\mathrm{PCA}}$ denote its mean score vector. At each step, SigDoTS merges the eligible pair with the smallest linkage value,

$$\Delta(A, B) = \frac{n_A n_B}{n_A + n_B} \left\| \overline{Z}_A^{\mathrm{PCA}} - \overline{Z}_B^{\mathrm{PCA}} \right\|_2^2 . \tag{S18}$$

Starting from the $K_0$ leaf domains, repeated merging yields a single root when the leaf-domain adjacency graph is connected. The resulting binary tree $\mathcal{T}^{\mathrm{obs}}$ contains unions of complete leaf domains, and the two children of each internal node define a candidate binary split.

**Stage 2: Nodewise two-domain test** At each visited internal node, SigDoTS compares the observed child split with the separation expected under a fitted node-specific single-mean null. Let $v$ denote an internal node of $\mathcal{T}^{\mathrm{obs}}$, with spatial-unit index set $I_v$, children $v_L$ and $v_R$, and size $n_v = |I_v|$. The two children define the observed assignment vector $\hat{z}_v^{\mathrm{obs}}$. On the node-restricted gene-expression matrix $X_v = X[I_v, \cdot]$, SigDoTS computes the observed separation statistic $\mathrm{CI}_{\mathrm{obs},v} = \mathrm{CI}(X_v, \hat{z}_v^{\mathrm{obs}})$ using the same gene-expression transformation and cluster-index definition as in the two-domain test.

SigDoTS fits the following null model within node $v$:

$$H_{0,v}: \quad X_v \sim \mathcal{MN}_{n_v \times p}\left(\mathbf{1}_{n_v} \mu_v^\top, \Sigma_{s,v}, \Sigma_{g,v}\right). \tag{S19}$$

Under $H_{0,v}$, all spatial units in the node share a common mean, while $\Sigma_{s,v}$ and $\Sigma_{g,v}$ describe node-specific spatial and gene covariance. SigDoTS estimates these parameters from $X_v$ and the node coordinates $S_v$ using the procedure defined for the two-domain test.

SigDoTS constructs the null reference distribution by repeating the complete node-local split-selection procedure in each replicate. Let $K_v$ denote the number of observed leaf domains descending from node $v$. SigDoTS holds $K_v$ fixed for the nodewise test because the domain-detection pipeline requires the number of initial candidate domains. For replicate $b$, SigDoTS simulates $\widetilde{X}_v^{(b)}$ from the fitted null and repeats the node-local $K_v$-domain detection and constrained tree construction on $\widetilde{X}_v^{(b)}$ and $S_v$. The two children of the replicate-specific root define $\widetilde{z}_v^{(b)}$, and the corresponding null statistic is $\widetilde{\mathrm{CI}}_v^{(b)} = \mathrm{CI}(\widetilde{X}_v^{(b)}, \widetilde{z}_v^{(b)})$. Each null statistic therefore incorporates both $K_v$-domain detection and constrained aggregation.

Smaller cluster-index values represent greater separation. SigDoTS computes the lower-tail Monte Carlo parametric-bootstrap $p$-value

$$\widehat{p}_v = \frac{1 + \sum_{b=1}^{B_v} \mathbf{1}\left\{\widetilde{\mathrm{CI}}_v^{(b)} \leq \mathrm{CI}_{\mathrm{obs},v}\right\}}{1 + B_v}, \tag{S20}$$

where $B_v$ is the number of null replicates generated for node $v$. SigDoTS also provides a normal-adaptive approximation of $\widehat{p}_v$ that requires fewer null replicates. It generates 10 null replicates and sets $\widehat{p}_v = \Phi\{(\mathrm{CI}_{\mathrm{obs},v} - \widehat{\mu}_v)/\widehat{\sigma}_v\}$, where $\widehat{\mu}_v$ and $\widehat{\sigma}_v$ are the sample mean and standard deviation of $\widetilde{\mathrm{CI}}_v^{(b)}$. When this value lies between $\alpha_v^\star/10$ and $10\alpha_v^\star$, with $\alpha_v^\star$ defined in Eq. (S21), SigDoTS increases the number of null replicates to 50 and recomputes $\widehat{p}_v$.

**Stage 3: Top-down hierarchical testing** SigDoTS tests the hierarchy from the root downward. The family-wise error rate (FWER) is the probability of rejecting at least one true nodewise null. SigDoTS tests node $v$ at

$$\alpha_v^\star = \alpha \frac{n_v - 1}{n - 1}, \tag{S21}$$

where $n_v$ and $n$ denote the numbers of spatial units in node $v$ and the full analyzed region. The root is tested at level $\alpha$, and the threshold decreases with node size.

If $\widehat{p}_v \le \alpha_v^\star$, SigDoTS rejects $H_{0,v}$, retains the observed split, and proceeds to child nodes that meet the prespecified size criteria. If $\widehat{p}_v > \alpha_v^\star$, SigDoTS fails to reject $H_{0,v}$, stops testing that branch, and retains $v$ as an unsplit terminal node. Leaves and nodes that do not meet the size criteria are also terminal. When the nodewise $p$-values are valid, testing at $\alpha_v^\star$ and stopping below each node that fails to reject $H_{0,v}$ control the FWER at level $\alpha$ [13, 14].

#### S1.2 Simulated studies

We generated gene count matrices $C = (C_{ig}) \in \mathbb{N}_0^{n\times p}$ for $n$ spatial units and $p$ genes using a Poisson log-normal model [15]. Specifically, given a rectangular region with $n$ spatial units arranged on a regular lattice, we first sampled a latent log-intensity field $Z \in \mathbb{R}^{n\times p}$ from a matrix-normal distribution,

$$Z \sim \mathcal{MN}_{n\times p}(M, \Sigma_s, \Sigma_g).$$

Here, $M = \mathbb{E}(Z) \in \mathbb{R}^{n\times p}$ was the mean field of the latent log-intensities, and the matrix-normal model implied $\mathrm{Cov}\{\mathrm{vec}(Z)\} = \Sigma_g \otimes \Sigma_s$. We independently sampled sequencing depth $L_i \sim \mathrm{Gamma}(\mathrm{shape} = 10, \mathrm{scale} = 500)$ for each spatial unit. Conditional on $Z$ and $L_i$, we sampled counts independently as

$$C_{ig} \mid Z_{ig}, L_i \sim \mathrm{Poisson}\{L_i \exp(Z_{ig})\}.$$

Thus, $\mathbb{E}(C_{ig} \mid Z_{ig}, L_i) = L_i \exp(Z_{ig})$, and $Z_{ig}$ determined the gene-specific latent log-intensity, whereas $L_i$ scaled this intensity by the library-size factor for spatial unit $i$.

For spatial units $i$ and $j$ separated by Euclidean distance $d_{ij}$, we defined the Matérn-3/2 spatial covariance as

$$(\Sigma_s)_{ij} = \left(1 + \frac{\sqrt{3}d_{ij}}{\ell}\right) \exp\left(-\frac{\sqrt{3}d_{ij}}{\ell}\right),$$

To match the spatial dependence to the local lattice spacing without using expression values, we set $\ell$ to the median fifth-nearest-neighbor distance. To introduce fixed low-rank gene–gene dependence, we used a spiked covariance model [16], setting

$$\Sigma_g = 0.7 I_p + 1.8 UU^\top,$$

where $U \in \mathbb{R}^{p\times 20}$ had random orthonormal columns. This covariance had eigenvalues 2.5 along the 20 factor directions and 0.7 along the remaining directions.

We considered five simulation scenarios:

- First, under the global null, all spatial units shared a latent log-rate mean of zero for every gene. We generated 100 replicate datasets at each of two sizes—144 and 2,000 spatial units—with 2,000 genes per dataset, to assess how frequently each method inferred multiple spatial domains in the absence of domain-specific mean shifts.
- Second, under balanced alternatives, we simulated three equal-width vertical domains. All genes had a baseline latent log-rate mean of zero. For 200 randomly selected genes, we shifted this mean to $-\delta/2$, $+\delta/2$ and $+2\delta$ across the three domains; the remaining genes retained a mean of zero. We added mean-zero spatial noise around these latent log-rate means. Thus, $\delta$ controls the separation between domain means on the latent log-rate scale. We generated 40 replicate datasets for the main setting with $\delta = 1.3$. To assess sensitivity to signal strength, we varied $\delta$ from 0.9 to 1.7, using 10 replicates per setting. We also generated 40 replicate datasets with five equal-width vertical domains, on a $20 \times 20$ lattice (400 spatial units) and a $50 \times 40$ lattice (2,000 spatial units), with latent log-rate mean shifts of $-2\delta$, $-\delta$, 0, $+\delta$ and $+2\delta$ relative to the same zero baseline.
- Third, under the unbalanced alternative, we retained the three-domain mean structure but varied the first domain from 24 to 144 spatial units while fixing each of the other two domains at 48 spatial units. We generated 40 replicate datasets at each of the six domain-size settings.

- Fourth, we varied the strength and the range of the spatial dependence. We generated these datasets on a $50 \times 50$ lattice of 2,500 spatial units with the spatial covariance

$$(\Sigma_s)_{ij} = \eta K_\ell(s_i, s_j) + (1 - \eta)\mathbf{1}\{i = j\},$$

  This is the spatial covariance model that SigDoTS fits (Section S1.1.3). Here, $K_\ell$ is the Matérn-3/2 kernel defined above, $\eta \in [0, 1]$ is the fraction of the latent variance that is spatially correlated, and $\ell$ is the length scale in lattice spacings. We used 17 settings: $\eta = 0$, and every combination of $\eta \in \{0.25, 0.5, 0.75, 1\}$ with $\ell \in \{\sqrt{2}, 3, 5, 10\}$. The spatial covariance of the first three scenarios corresponds to $\eta = 1$ and $\ell = \sqrt{2}$, because the median fifth-nearest-neighbor distance on a lattice is $\sqrt{2}$ lattice spacings. At each setting, we generated 20 global-null datasets and 20 three-domain datasets. The three domains were vertical bands of 850, 850 and 800 spatial units, with the mean shifts $-\delta/2$, $+\delta/2$ and $+2\delta$ of the balanced alternative and $\delta = 1.3$, applied to 1,000 randomly selected genes.
- Fifth, to assess computational cost, we also generated three-domain datasets on six square lattices of 400, 900, 1,600, 2,500, 5,041 and 10,000 spatial units, two datasets per lattice, at ($\eta = 1$) and ($\ell = \sqrt{2}$) and with the same mean shifts and (1,000) signal genes as above.

For each simulated dataset, we log-normalized counts per 10,000 and generated candidate domains using $k$-means on the first 15 principal components. We used $K_0 = 5$ for the null, three-domain and unbalanced settings and $K_0 = 8$ for the five-domain setting. We compared SigDoTS, HC-Wald and scSHC [12] using these shared initial candidate domains, with $\alpha = 0.05$. SigDoTS constructed the candidate-domain hierarchy in the same principal-component space and repeated the clustering procedure for every null replicate. scSHC used raw counts, all genes and 15 principal components. HC-Wald tested the same hierarchy and used the same nodewise thresholds $\alpha_v^\star$ as SigDoTS, treating candidate domains as prespecified and comparing the children's mean principal-component scores using a Wald test with a $\chi^2_{15}$ reference distribution.

In addition to the unbalanced alternative, we performed two sensitivity analyses in which we applied only SigDoTS. Both used the three-domain mean structure of the balanced alternative with $\delta = 1.3$, 2,000 genes and 300 signal genes, and generated 100 global-null and 10 three-domain datasets per setting. First, to assess dependence on the upstream domain-detection method, we generated $K_0 = 10$ initial candidate domains using BANKSY [17], DR-SC [18] or $k$-means. BANKSY used $\lambda = 0.2$ and $k$-means clustering of 20 BANKSY principal components, and DR-SC used 15 latent dimensions. The global-null datasets contained 144 spatial units on a $12 \times 12$ lattice to limit computational cost, and the three-domain datasets contained 2,000 spatial units on a $50 \times 40$ lattice. Second, to assess dependence on the initial candidate-domain count, we generated $K_0 \in \{4, 6, 9, 12, 18\}$ initial candidate domains using $k$-means on a $12 \times 12$ lattice. In both analyses, SigDoTS applied the same upstream method to every null replicate and computed nodewise $p$-values with the normal-adaptive approximation; we repeated both analyses with the Monte Carlo $p$-value in Eq. (S20) for comparison.

Under the null, we evaluated the root node's rejection rate and quantile–quantile plots of root $p$-values. Under the alternatives, we evaluated domain recovery using the adjusted Rand index (ARI; Section S1.7) between the true and estimated domain assignments and the number of retained domains. For the computational-cost datasets, we report the elapsed time of each method and its peak memory, the largest of one-second samples of the memory used by all processes of the run. In the sensitivity analyses, we also computed the boundary F1 score (Section S1.7) and an oracle ARI, defined as the highest ARI among the domain assignments obtained by merging initial candidate domains along the observed hierarchy. The oracle ARI bounds the ARI that SigDoTS can attain from that hierarchy. We reported root rejection rates with two-sided 95% Wilson confidence intervals. In the fourth scenario, error bars show exact 95% Clopper–Pearson intervals for proportions and percentile bootstrap 95% intervals for mean ARI (2,000 resamples). Representative spatial maps and hierarchical testing trees illustrated the relationship among the true domains, initial candidate domains and retained domains.

#### S1.3 Human DLPFC Visium data analysis

We analyzed human dorsolateral prefrontal cortex (DLPFC) Visium sections 151673, 151674, 151675 and 151676 from donor 3 in the spatialLIBD R package [19]. We performed log-CPM normalization and identified the top 2,000 highly variable genes. Manual layer labels served only to define analysis masks and interpret the retained domains. We evaluated SigDoTS using two complementary tasks.

First, we tested whether SigDoTS retained domains corresponding to known laminar and cell-type organization. We focused on Layer 6 because previous spatial and single-cell analyses of human DLPFC identified molecularly distinct subdomains within this histological layer [20]. We included all manually labeled Layer 6 spots from the four sections, generated initial BANKSY candidate domains and applied SigDoTS to test the candidate splits. We used HC-Wald and scSHC as baselines (Section S1.2). For each section, we visualized the manual layer annotations, BANKSY candidate domains and domains retained by SigDoTS, HC-Wald and scSHC and their hierarchical testing trees. We designated the retained domains as superficial or deep Layer 6 according to their proximity to Layer 5 and white matter. We then evaluated predefined neuronal, glial and laminar marker programs. Dot plots summarized marker expression across domains and sections, and spatial maps localized representative markers. For the cortical-depth profiles, we normalized depth from 0 at the Layer 5-facing edge of Layer 6 to 1 at the white-matter-facing edge. Within each section, we calculated neuronal and myelination module scores by averaging the standardized expression of their predefined marker genes, and then averaged these scores within ten equal-width depth bins. These analyses assessed whether the retained split corresponded to established Layer 6 organization.

Second, we tested SigDoTS' ability to merge visually plausible candidate domains within a region with limited evidence of discrete molecular substructure. We selected an interior band of Layer 3 based on prior anatomical and molecular evidence and evaluated whether SigDoTS would retain a split imposed within a single annotated layer. Specifically, we restricted manually labeled Layer 3 spots to a cortical-depth interval of 0.25–0.75 to further restrict the analysis to the inner Layer 3 region. Within the inner Layer 3 region, we used BANKSY with $\lambda = 0.2$ to generate $K_0 = 2$ candidate domains and then applied SigDoTS to these candidate domains with $\alpha = 0.05$. We repeated the analysis with $K_0 = 2, 3, 4$ to determine whether SigDoTS rejected the single-domain null as the initial candidate-domain count increased. We also applied HC-Wald and scSHC with default settings as baselines.

To assess cell-type compositional homogeneity independently of SigDoTS, we used cell2location for cell type deconvolution. The reference was DLPFC single-nucleus RNA-sequencing data from the spatialDLPFC atlas [20], accessed through spatialLIBD. We estimated posterior mean cell-type abundances using the 26-category `cellType_hc` annotation, retained cell types with a mean abundance share of at least 0.01 within the Layer 3 interior and normalized the retained abundances to spot-level proportions. Spots with zero total retained abundance were excluded. We calculated local homogeneity $H_\ell$ over the unique undirected edges of a six-nearest-neighbor graph and global homogeneity $H_g$ across all spots in the Layer 3 interior (Section S1.7). Both scores range from 0 to 1. Values of $H_\ell$ near 1 indicate that neighboring spots have similar cell-type compositions, whereas values of $H_g$ near 1 indicate compositional similarity across the complete Layer 3 interior.

#### S1.4 Ovarian cancer Visium HD data analysis

We analyzed a SPATCH ovarian cancer FFPE Visium HD dataset at 8 $\mu$m bin resolution. We selected a region of interest (ROI) containing 19,192 in-tissue bins. Quality control retained 13,690 bins with at least 100 UMIs and 50 detected genes, and genes detected in at least 10 bins. We log-normalized counts per 10,000 and selected 2,000 highly variable genes using residual variance after mean–variance trend fitting. We standardized each selected gene across bins and then used BANKSY to obtain $K_0 = 15$ candidate spatial domains. Given the BANKSY candidate domains, we applied SigDoTS with hierarchical adjustment at an overall error level of $\alpha = 0.05$. Spatial covariance was modeled using a Matérn kernel with smoothness parameter $\nu = 3/2$.

To assess whether the spatial domains retained by SigDoTS differed in cell-type composition, we first used SPATCH transcriptomic labels transferred from matched single-cell RNA-sequencing data [21]. We matched these labels to Visium HD bins by bin index and quantified cell-type enrichment as the $\log_2$ ratio of each cell type's proportion within a domain to its ROI-wide proportion. For predefined RNA markers of epithelial, myeloid, fibroblast, CD8 T-cell, CD4 T-cell, plasma-cell, smooth-muscle and endothelial states, we calculated mean standardized expression and detection frequency within each domain.

We also assessed domain composition using manually annotated CODEX cells from an adjacent serial tissue section [21]. After cross-section registration, we mapped each CODEX cell centroid to its nearest analyzed bin within 12 $\mu$m and assigned each mapped cell the SigDoTS and BANKSY domain assignments of that bin. We calculated domain-specific cell-type proportions and mean protein intensities. For selected proteins, we expressed domain-specific mean intensities as fold changes relative to the mean across the other domains and mapped their spatial intensities across the ROI.

#### S1.5 Ovarian cancer Visium data analysis

We analyzed a 10x Genomics human ovarian tumor Visium dataset [22] containing $3,455$ spatial spots. After log-CPM normalization, we selected $2,000$ highly variable genes. We used BANKSY to obtain $K_0 = 15$ candidate spatial domains and applied SigDoTS with hierarchical adjustment at an overall error level of $\alpha = 0.05$, as in the Visium HD analysis above.

For visualization, we first displayed the BANKSY candidate domains alongside the final domains retained by SigDoTS. We then selected 27 marker genes corresponding to nine cell types. For each marker, we calculated the mean $z$-score within each retained domain together with the proportion of spots expressing that gene. We further selected one representative marker for each cell type and transformed the mean $z$-score to a relative marker score, $2^z$, to compare marker expression across retained domains. Finally, the spatial expression patterns of these nine representative markers were compared with the retained-domain assignments. We additionally examined representative candidate splits for which SigDoTS failed to reject the single-domain null by comparing their constituent BANKSY domains using the same cell-type and marker-expression summaries.

#### S1.6 Colorectal adenocarcinoma Xenium 5K data analysis

We analyzed colorectal adenocarcinoma (COAD) Xenium 5K data obtained from SPATCH, using gene-expression counts aggregated over segmented cells [21]. We selected a fixed rectangular ROI in the middle-right portion of the tissue section containing 25,343 cells flagged as high quality by SPATCH. We applied log-CPM normalization, selected 2,000 highly variable genes using residual variance after LOESS mean–variance trend fitting and standardized each selected gene across cells. We then used BANKSY to obtain $K_0 = 15$ candidate spatial domains. SigDoTS constructed a hierarchy over these domains using 20 principal components and tested candidate splits using the standardized gene-expression matrix, with hierarchical adjustment at an overall error level of $\alpha = 0.05$. Spatial covariance was modeled using a Matérn kernel with smoothness parameter $\nu = 3/2$.

We assessed domain cell-type composition using SPATCH transcriptomic annotations and expressed enrichment as the $\log_2$ ratio of observed to expected cell-type proportions. For predefined RNA markers of proliferation, plasma-cell, B-cell, epithelial, tumor-mucin, myeloid, mural and endothelial states, we calculated mean standardized expression within each domain.

We also analyzed manually annotated CODEX cells from an adjacent serial tissue section [21]. We matched CODEX cells to Xenium cells using shared identifiers when available and nearest-neighbor assignment within four pixels in the shared coordinate system otherwise. CODEX protein intensities were summarized and visualized as described for the Visium HD analysis above. We additionally mapped CODEX epithelial cells and B cells and calculated the percentage of all mapped B cells assigned to each domain retained by SigDoTS.

#### S1.7 Evaluation metrics

- **Adjusted Rand index.** For a true domain assignment $\mathcal{U} = \{U_1, \ldots, U_R\}$ and an estimated domain assignment $\mathcal{V} = \{V_1, \ldots, V_S\}$ of $n$ spatial units, let $n_{rs} = |U_r \cap V_s|$, $a_r = \sum_s n_{rs}$ and $b_s = \sum_r n_{rs}$. We defined the adjusted Rand index (ARI) as

$$\mathrm{ARI} = \frac{\displaystyle\sum_{r,s} \binom{n_{rs}}{2} - \frac{\sum_r \binom{a_r}{2} \sum_s \binom{b_s}{2}}{\binom{n}{2}}}{\displaystyle\frac{1}{2}\left\{\sum_r \binom{a_r}{2} + \sum_s \binom{b_s}{2}\right\} - \frac{\sum_r \binom{a_r}{2} \sum_s \binom{b_s}{2}}{\binom{n}{2}}}.$$

  The ARI equals 1 for identical domain assignments and has expected value 0 under random assignment; negative values indicate less agreement than expected by chance.
- **Boundary F1 score.** We built a six-nearest-neighbor graph on the spatial units and defined an edge as a boundary edge when its two endpoints had different domain assignments. Let $E_{\mathrm{true}}$ and $E_{\mathrm{est}}$ denote the boundary edges under the true and estimated domain assignments. With precision $P = |E_{\mathrm{true}} \cap E_{\mathrm{est}}|/|E_{\mathrm{est}}|$ and recall $R = |E_{\mathrm{true}} \cap E_{\mathrm{est}}|/|E_{\mathrm{true}}|$, we defined the boundary F1 score as $2PR/(P + R)$. The score equals 1 when the estimated and true boundaries coincide.
- **Normalized cell-type composition.** For a region containing $n$ spatial units and $C$ cell types, let $a_{ik}$ denote the estimated abundance of cell type $k$ at spatial unit $i$. Using the same set of cell types across all spatial units, we defined the normalized composition

$$p_{ik} = \frac{a_{ik}}{\sum_{k'=1}^{C} a_{ik'}}, \qquad p_i = (p_{i1}, \ldots, p_{iC}).$$

  Spatial units with zero total abundance were excluded.

- **Local Jensen–Shannon homogeneity.** For spatial units $i$ and $j$, let $m_{ij} = (p_i + p_j)/2$ and define

$$\mathrm{JSD}_2(p_i, p_j) = \frac{1}{2}\sum_{k=1}^{C} p_{ik}\log_2\frac{p_{ik}}{m_{ij,k}} + \frac{1}{2}\sum_{k=1}^{C} p_{jk}\log_2\frac{p_{jk}}{m_{ij,k}},$$

  with $0\log_2 0 = 0$. Let $E$ be the set of unique undirected edges in a prespecified spatial-neighbor graph. We defined local homogeneity as

$$H_\ell = 1 - \frac{1}{|E|}\sum_{(i,j)\in E}\mathrm{JSD}_2(p_i, p_j).$$

  The score satisfies $0 \le H_\ell \le 1$, with values near 1 indicating similar compositions among neighboring spatial units. A smooth region-wide gradient can produce a high $H_\ell$ because adjacent spatial units may remain similar.
- **Global Jensen–Shannon homogeneity.** Let

$$\bar{p} = \frac{1}{n}\sum_{i=1}^{n} p_i, \qquad h_2(p) = -\sum_{k=1}^{C} p_k \log_2 p_k.$$

  We defined global homogeneity as

$$H_g = 1 - \frac{h_2(\bar{p}) - n^{-1}\sum_{i=1}^{n} h_2(p_i)}{\log_2\{\min(n, C)\}}.$$

  The score satisfies $0 \le H_g \le 1$, with values near 1 indicating similar compositions across the complete region. Unlike $H_\ell$, $H_g$ captures broad gradients or spatially coherent patches that differ in composition.
- **Domain-wise cell-type enrichment.** For domain $d$, let $I_d$ denote its spatial-unit index set and $n_d = |I_d|$. We defined the domain and region-wide mean proportions of cell type $k$ as

$$\bar{p}_{dk} = \frac{1}{n_d}\sum_{i\in I_d} p_{ik}, \qquad \bar{p}_{\cdot k} = \frac{1}{n}\sum_{i=1}^{n} p_{ik}.$$

  The domain-wise enrichment score was

$$R_{dk} = \log_2\left(\frac{\bar{p}_{dk} + \varepsilon}{\bar{p}_{\cdot k} + \varepsilon}\right), \qquad \varepsilon = 10^{-3}.$$

  A score of $R_{dk} = 0$ indicates that cell type $k$ has the same mean proportion in domain $d$ as in the complete region; positive and negative values indicate enrichment and depletion, respectively. This score is descriptive and does not constitute a hypothesis test.

### S2 Supplementary Results

#### S2.1 Additional simulation results

This section gives details of the sensitivity analyses summarized in the main text (Supplementary Figs. S1 and S2).

Across upstream domain-detection methods, BANKSY, DR-SC and $k$-means each generated $K_0 = 10$ initial candidate domains. Under the global null, root rejection rates were at or below $\alpha = 0.05$, SigDoTS retained a median of one domain for each method, and the empirical distribution of root $p$-values was approximately uniform on $[0, 1]$ (Supplementary Fig. S2a,b). Under the three-domain alternative, SigDoTS retained a median of three domains, with median ARI and boundary F1 scores of 1.000 for each upstream method (Supplementary Fig. S2c–e). Its median ARI equaled the oracle benchmark, the highest ARI obtainable by merging initial candidate domains along each observed hierarchy.

Across initial candidate-domain counts $K_0 \in \{4, 6, 9, 12, 18\}$, SigDoTS retained a median of one domain under the global null at every $K_0$, and the empirical distribution of root $p$-values remained approximately uniform on $[0, 1]$ (Supplementary Fig. S2f,g). Under the three-domain alternative, the ARI and boundary F1 scores of the initial candidate domains decreased as $K_0$ increased. SigDoTS improved both measures at every $K_0$ while retaining a median of three domains (Supplementary Fig. S2h). Both measures nevertheless decreased at $K_0 = 18$, the largest $K_0$ tested.

In the domain-size imbalance analysis, the first domain varied from 2 to 12 columns (24 to 144 spatial units) while the other two domains had 4 columns (48 spatial units) each (Supplementary Fig. S2i–k). For first-domain sizes of 48–144 spatial units, SigDoTS retained three domains in 37 to 40 of 40 datasets per setting and achieved a median ARI of 1.000, recovering the true domains even when one domain contained three times as many spatial units as the others. HC-Wald retained a median of five domains at every size. As the first domain increased from 48 to 144 spatial units, scSHC retained more domains and its median ARI decreased. At the two largest sizes, both baselines retained a median of five domains, indicating oversegmentation despite the increase in the number of spatial units (Supplementary Fig. S2l). At 24 spatial units, SigDoTS retained two domains in 36 of 40 datasets and achieved a lower median ARI than scSHC but a higher median ARI than HC-Wald.

Across signal strengths with 2,000 spatial units, SigDoTS retained $K = 3$ in all 10 datasets for $\delta \leq 1.5$ and in 9 of 10 datasets at $\delta = 1.7$ (Supplementary Fig. S1d). By contrast, HC-Wald retained all five initial candidate domains in every dataset at both sizes, whereas scSHC retained a mean of 3.4 to 3.9 domains with 144 spatial units and all five with 2,000 spatial units.

#### S2.2 Additional results for the datasets

In the Layer 3 interior of DLPFC section 151673, HC-Wald rejected the single-domain null at every tested node ($p < 0.001$) for $K_0 = 2$, 3 and 4. scSHC retained one domain for $K_0 = 2$ and $K_0 = 4$ (root $p = 0.980$ and 0.890) but two domains for $K_0 = 3$, for which its root test rejected the single-domain null ($p < 0.001$) whereas the subsequent test did not ($p = 1.000$).

In the ovarian cancer Visium HD data, HLA-DR can be expressed by antigen-presenting cells, activated T cells and ovarian tumor epithelial cells [23, 24]; we therefore interpreted D1 as a tumor-epithelial domain and D2 as an epithelial domain with stronger T-cell- and MHC class II-associated signals. The distinction between the fibroblast-containing domains D4 and D5 is consistent with spatial profiling of HGSOC showing distinct CAF-associated plasma-cell compartments and variation in T-cell localization [25]. Within D6, adjacent-section CODEX localized the largest difference to candidate domain 8, which had stronger CD34 signal and a higher endothelial-cell fraction than candidate domains 1 and 7 (Supplementary Fig. S5b,c). Cell-type composition differed among the three candidate domains ($p = 0.0128$; Cramér's $V = 0.302$), but after BH correction, only candidate domain 8 differed from the other two ($p = 0.0043$, $q = 0.0129$; Cramér's $V = 0.360$; Supplementary Fig. S5d). In the ovarian cancer Visium data, epithelial markers PAX8 and EPCAM were highest in D9, fibroblast markers DCN and COL1A1 were higher in D2 and D5–D7, and endothelial markers PECAM1 and VWF were highest in D4 (Supplementary Fig. S6d). Relative marker scores summarized these domain-level differences, and spatial expression maps localized them to different tissue regions (Supplementary Fig. S6f,g). As in Visium HD, the fibroblast-associated domains differed in their accompanying immune signals: D2 had higher expression of T-cell markers CD3D, CD3E and TRBC1 and plasma-cell markers MZB1 and JCHAIN than D5–D7 (Supplementary Fig. S6d). SigDoTS merged seven BANKSY candidate domains into the epithelial domain D9 (Supplementary Fig. S6h); the single-domain null was not rejected at their common parent node ($p = 0.218$; $p_{\text{adj}} = 0.446$; Supplementary Fig. S6e), and the candidate domains shared PAX8 and EPCAM expression

(Supplementary Fig. S6i).

In the colorectal adenocarcinoma Xenium data, the separation of epithelial domains D1, D4 and D5 is in line with previous single-cell studies that identified distinct epithelial differentiation and intrinsic transcriptional states in colorectal tumors [26, 27]. Within D6, spatially separated T-cell-rich and macrophage-rich niches have also been reported in colorectal cancer [28]. Adjacent-section CODEX cell-type composition differed across the five candidate domains (Cramér's $V = 0.252$, $p = 0.0001$; Supplementary Fig. S8b–d). Candidate domain 7 had the largest one-versus-rest compositional difference (Cramér's $V = 0.468$, BH-adjusted $q = 0.000250$), the highest CD4-T-cell fraction and higher CD3e, CD4, HLA-A and HLA-DR protein signals than the other candidate domains. Within D7, CODEX cell-type composition differed across candidate domains 2, 14, 13 and 1 (Cramér's $V = 0.322$, $p = 0.0001$), with candidate domain 13 having the largest fibroblast fraction and SMA protein signal (Supplementary Fig. S9b–d). D7 therefore contained localized fibroblast-associated variation, but its candidate domains were not retained separately after hierarchical adjustment.

#### S2.3 Reduced computational cost with the normal-adaptive implementation

To reduce the computational cost of null calibration, we developed a normal-adaptive implementation of SigDoTS that uses a normal approximation and adds null replicates for tests near the rejection threshold (Section S1.1.6). It produced ARI values and retained domain counts similar to those of the Monte Carlo bootstrap across the three- and five-domain alternatives, domain sizes and signal strengths (Supplementary Fig. S10a–d). Both implementations had global-null root rejection rates below the nominal level of $\alpha = 0.05$ in these simulations (Supplementary Fig. S10e). Agreement in domain recovery also extended across initial domain counts (Supplementary Fig. S11a–e) and upstream domain-detection methods (Supplementary Fig. S12a–f), with the Monte Carlo bootstrap producing more conservative null rejection rates.

### S3 Data Availability

All experimental datasets analyzed in this study are publicly available. The human DLPFC Visium data for sections 151673, 151674, 151675 and 151676, including spatial coordinates and manual cortical-layer annotations, are available through spatialLIBD (https://research.libd.org/spatialLIBD/) [19]. The single-nucleus RNA-sequencing reference used for cell-type deconvolution is available from the spatialDLPFC atlas (https://research.libd.org/spatialDLPFC/) and can also be accessed through spatialLIBD [20]. The human ovarian cancer Visium dataset is available from 10x Genomics (https://www.10xgenomics.com/datasets/human-ovarian-cancer-1-standard) [22]. The ovarian cancer FFPE Visium HD and colorectal adenocarcinoma Xenium 5K datasets, together with the associated transcriptomic cell-type annotations and adjacent-section CODEX data, are available from SPATCH (https://spatch.pku-genomics.org/) [21].

### S4 Supplementary Figures

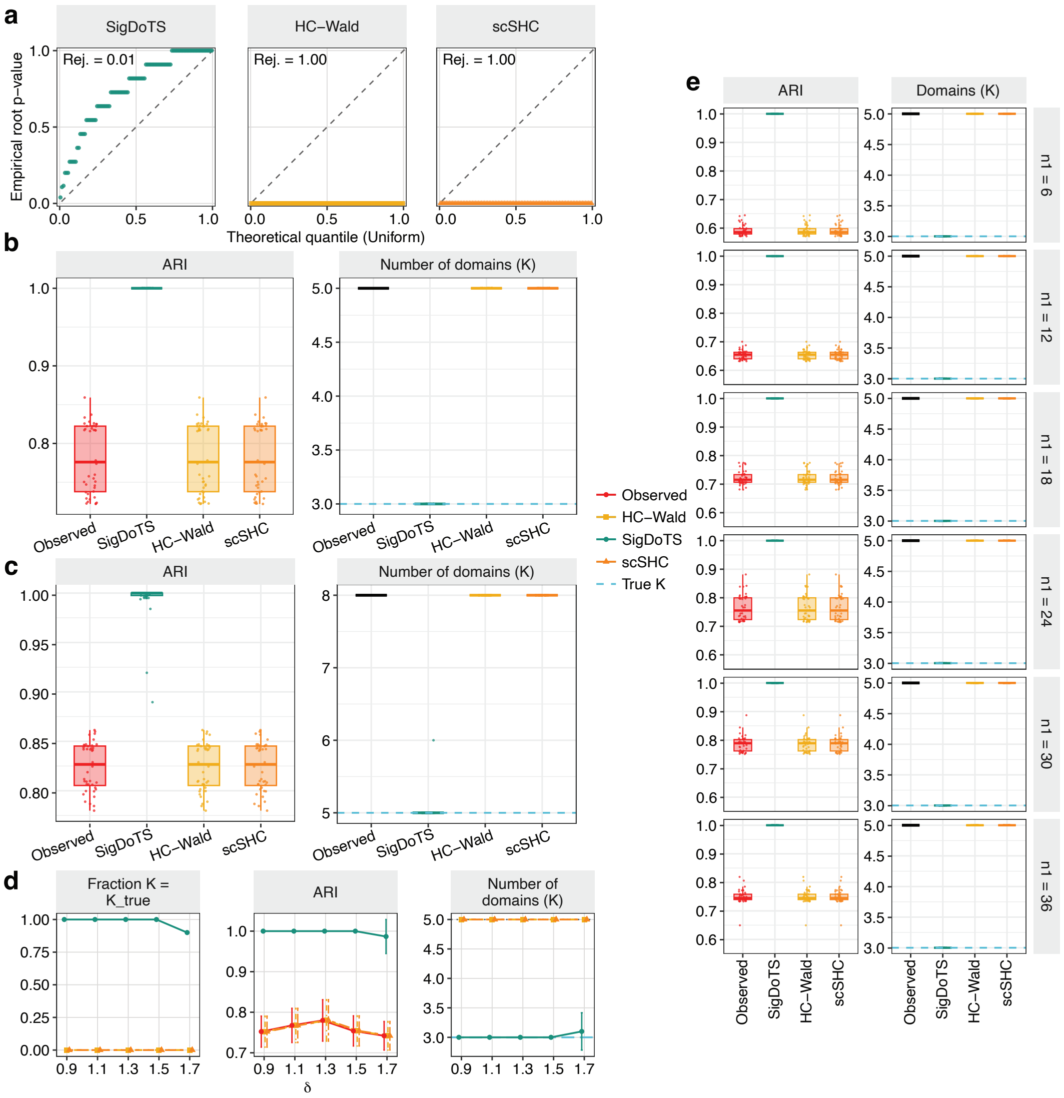

**Supplementary Fig. S1: Domain recovery and null calibration in simulations with a baseline size of 2,000 spatial units.** SigDoTS used the Monte Carlo bootstrap. **a** Root $p$-value quantile–quantile plots for 100 global-null datasets. "Rej." denotes the root rejection rate at $\alpha = 0.05$; diagonals mark the uniform reference. **b** Adjusted Rand index (ARI; left) and number of retained domains $K$ (right) for the three-domain alternative ($K_{\text{true}} = 3$, initial candidate-domain count $K_0 = 5$; 40 replicates). "Observed" denotes the initial candidate domains. The blue dashed line marks $K_{\text{true}}$; the black bar marks $K_0$. Points denote individual datasets; boxplots show medians and interquartile ranges, with whiskers reaching the most extreme values within 1.5× the interquartile range of the box edges. **c** ARI and retained $K$ for the five-domain alternative ($K_{\text{true}} = 5$, $K_0 = 8$; 40 replicates). Plotting conventions follow **b**. **d** Proportion of datasets with $K = K_{\text{true}}$, mean ARI and mean retained $K$ as the mean-difference parameter $\delta$ varied from 0.9 to 1.7 (10 replicates per setting). Larger $\delta$ indicates greater differences between domain means. Error bars show ±1 s.d. for ARI and $K$; blue dashed and gray dotted lines mark $K_{\text{true}} = 3$ and $K_0 = 5$, respectively. **e** ARI (left) and retained $K$ (right) as domain D1 varied from $n_1 = 6$ to 36 columns, with 16 columns in each remaining domain and 40 rows ($\delta = 1.3$; 40 replicates per setting). Plotting conventions follow **b**, with $K_{\text{true}} = 3$ and $K_0 = 5$.

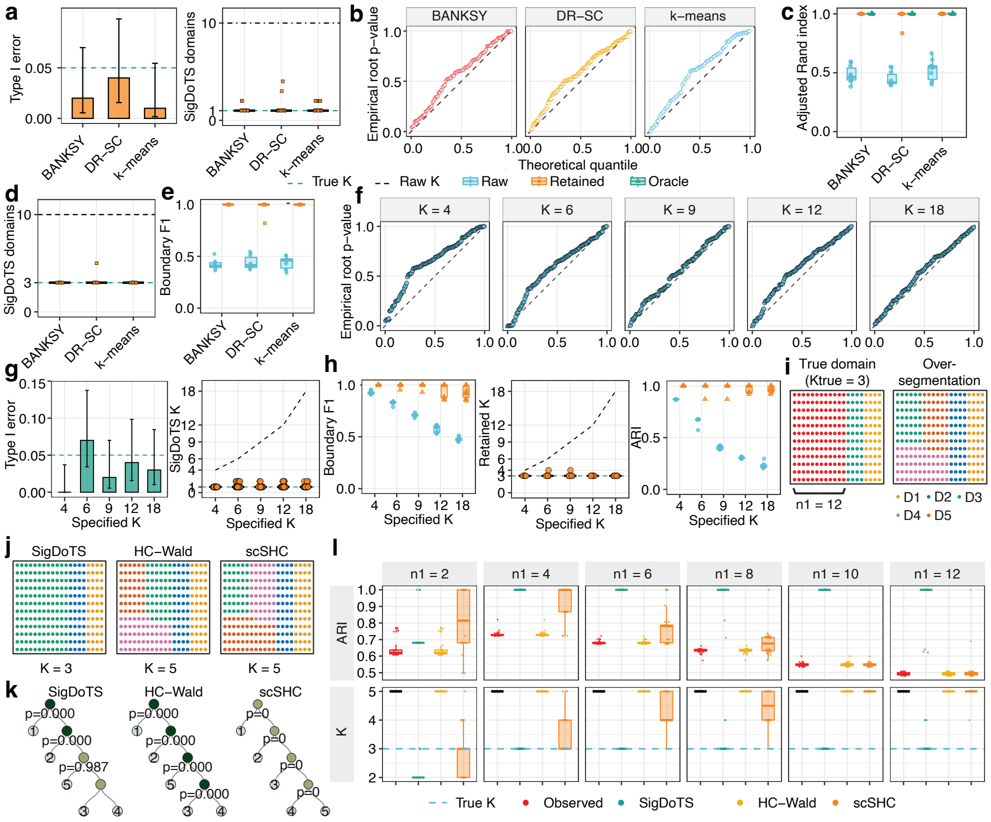

**Supplementary Fig. S2: Sensitivity of SigDoTS to upstream methods, initial candidate-domain counts and domain-size imbalance.** **a** Empirical type I error (left) and number of retained domains $K$ (right) under the global null for BANKSY, DR-SC and $k$-means initial candidate domains ($K_0 = 10$). Bars show the proportion of null datasets with root rejection; error bars show two-sided 95% Wilson confidence intervals, and the dashed line marks $\alpha = 0.05$. Reference lines mark $K_{\text{true}} = 1$ and $K_0 = 10$. Points denote individual datasets; boxes show medians and interquartile ranges, with whiskers reaching the most extreme values within 1.5× the interquartile range of the box edges. **b** Root $p$-value quantile–quantile plots under the global null for the upstream methods in **a**. Diagonals mark the uniform reference. **c** Adjusted Rand index (ARI) under the three-domain alternative with $n = 2000$ spatial units and $K_0 = 10$. "Raw" denotes the initial candidate domains; "Oracle" denotes merging initial candidate domains along the hierarchy to maximize ARI against the true domain labels, providing an upper bound on domain recovery achievable within that hierarchy. Points and boxplots are defined as in **a**. **d** Number of retained domains $K$ under the three-domain alternative. Reference lines mark $K_{\text{true}} = 3$ and $K_0 = 10$; points and boxplots are defined as in **a**. **e** Boundary F1 under the three-domain alternative. Points and boxplots are defined as in **a**. **f** Root $p$-value quantile–quantile plots under the global null as $k$-means generated $K_0 \in \{4, 6, 9, 12, 18\}$ initial candidate domains; diagonals mark the uniform reference. **g** Global-null root rejection rate (left) and retained $K$ (right) across the $K_0$ settings in **f**. Bars and error bars are defined as in **a**, and the dashed line marks $\alpha = 0.05$. Domain-number reference lines mark $K_{\text{true}} = 1$ and $K_0$; points and boxplots are defined as in **a**. **h** Boundary F1 (left), ARI (middle) and retained $K$ (right) under the three-domain alternative across the $K_0$ settings in **f**. "Raw" denotes the initial candidate domains. Points and boxplots are defined as in **a**; reference lines mark $K_{\text{true}} = 3$ and $K_0$. **i** Representative unbalanced three-domain alternative with $n_1 = 12$ columns in domain 1 and four in each remaining domain, together with its $K_0 = 5$ initial candidate domains. **j** Domains retained by SigDoTS, HC-Wald and scSHC for the dataset in **i**. **k** Testing trees for the dataset in **i**. Internal-node labels show $p$-values; leaf labels identify initial candidate domains. **l** ARI and retained $K$ as domain 1 varied over $n_1 \in \{2, 4, 6, 8, 10, 12\}$ columns, with four columns in each remaining domain and 12 rows (40 datasets per setting). "Observed" denotes the initial candidate domains. Points and boxplots are defined as in **a**; the dashed line marks $K_{\text{true}} = 3$.

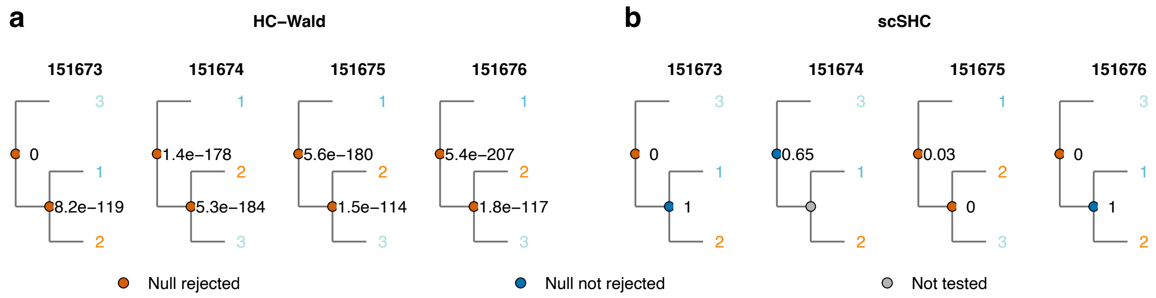

**Supplementary Fig. S3: Hierarchical testing of DLPFC Layer 6 by HC-Wald and scSHC.** Testing trees for HC-Wald (**a**) and scSHC (**b**) in human dorsolateral prefrontal cortex (DLPFC) sections 151673, 151674, 151675 and 151676, starting from three initial BANKSY candidate domains per section. Node labels give $p$-values, and leaf labels identify initial candidate domains.

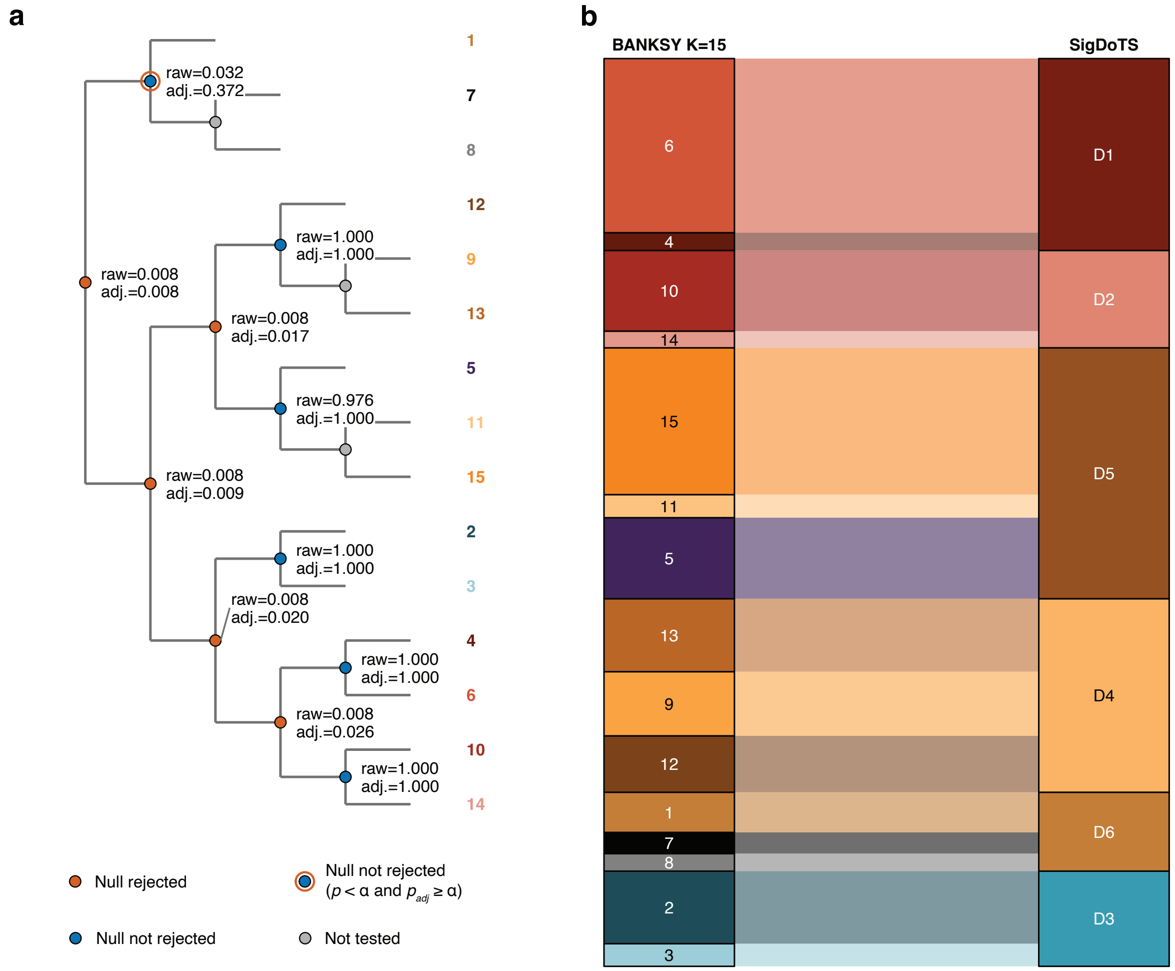

**Supplementary Fig. S4: Hierarchical testing and domain assignment in the ovarian cancer Visium HD dataset. a** Hierarchical testing tree for the 15 BANKSY candidate domains. Labels at tested nodes show the unadjusted $p$-value and the hierarchically adjusted $p_{\text{adj}}$-value. Red nodes denote rejection after hierarchical adjustment; blue nodes denote failure to reject before adjustment; red-outlined blue nodes denote $p < \alpha$ and $p_{\text{adj}} \geq \alpha$; and gray nodes denote tests not performed under the hierarchical stopping rule. Leaf-label colors indicate the corresponding retained-domain assignments. **b** Mapping of the 15 BANKSY candidate domains ($K_0 = 15$) to the six retained domains, D1–D6.

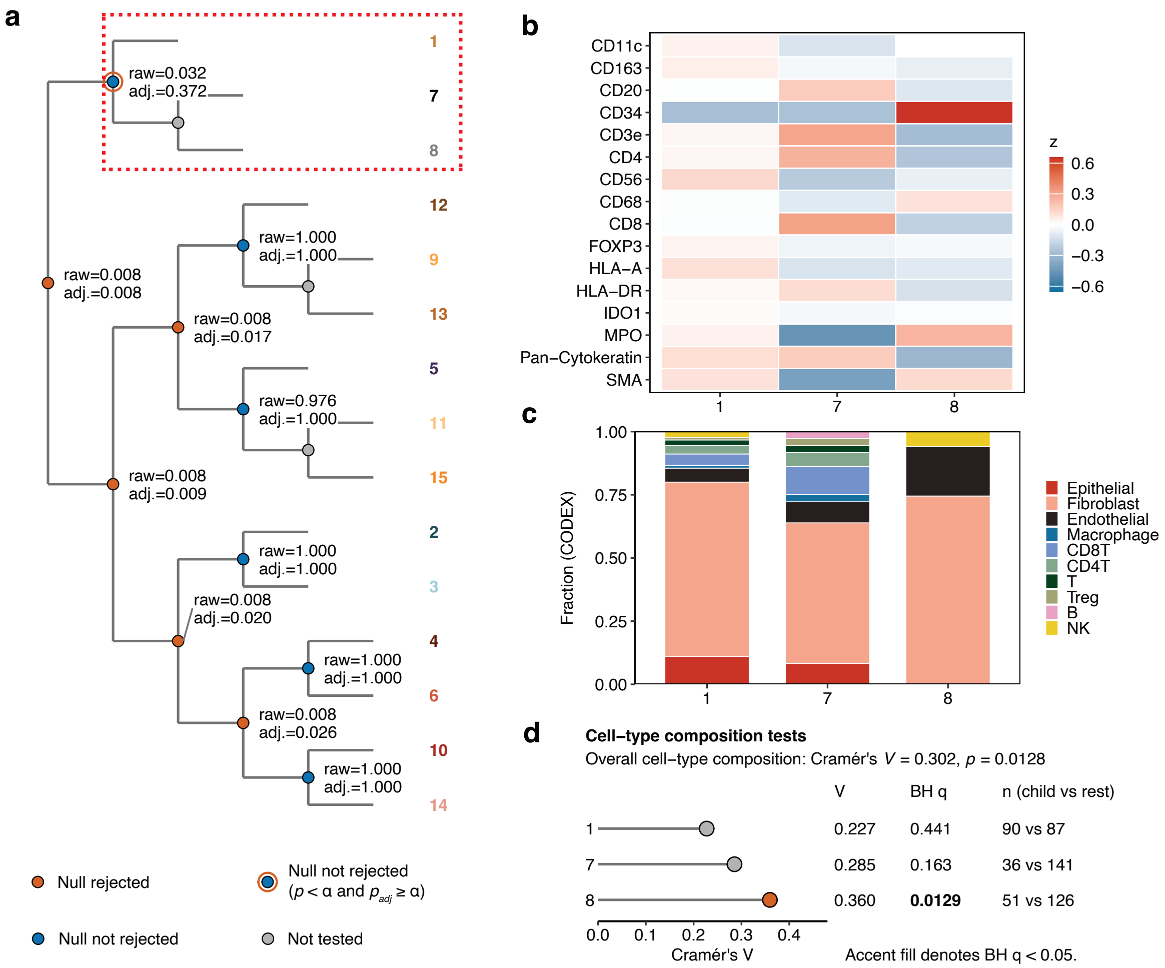

**Supplementary Fig. S5: CODEX cell-type composition differs among candidate domains merged into D6 in the ovarian cancer Visium HD dataset.** **a** Hierarchical testing tree with the D6 candidate split outlined in red. **b** Mean CODEX protein intensities, displayed as marker-wise $z$-scores, for adjacent-section cells assigned to candidate domains 1, 7 and 8. **c** CODEX cell-type fractions within the three candidate domains. **d** Overall and candidate-domain-versus-rest analyses of CODEX cell-type composition. Points show Cramér's $V$ for each candidate domain compared with the remaining candidate domains pooled. The table reports Benjamini–Hochberg (BH)-adjusted $q$ values and the corresponding CODEX cell counts. Accent-filled points denote BH-adjusted $q < 0.05$. CODEX measurements were obtained from the adjacent section and were not used for candidate-domain detection or hierarchical testing.

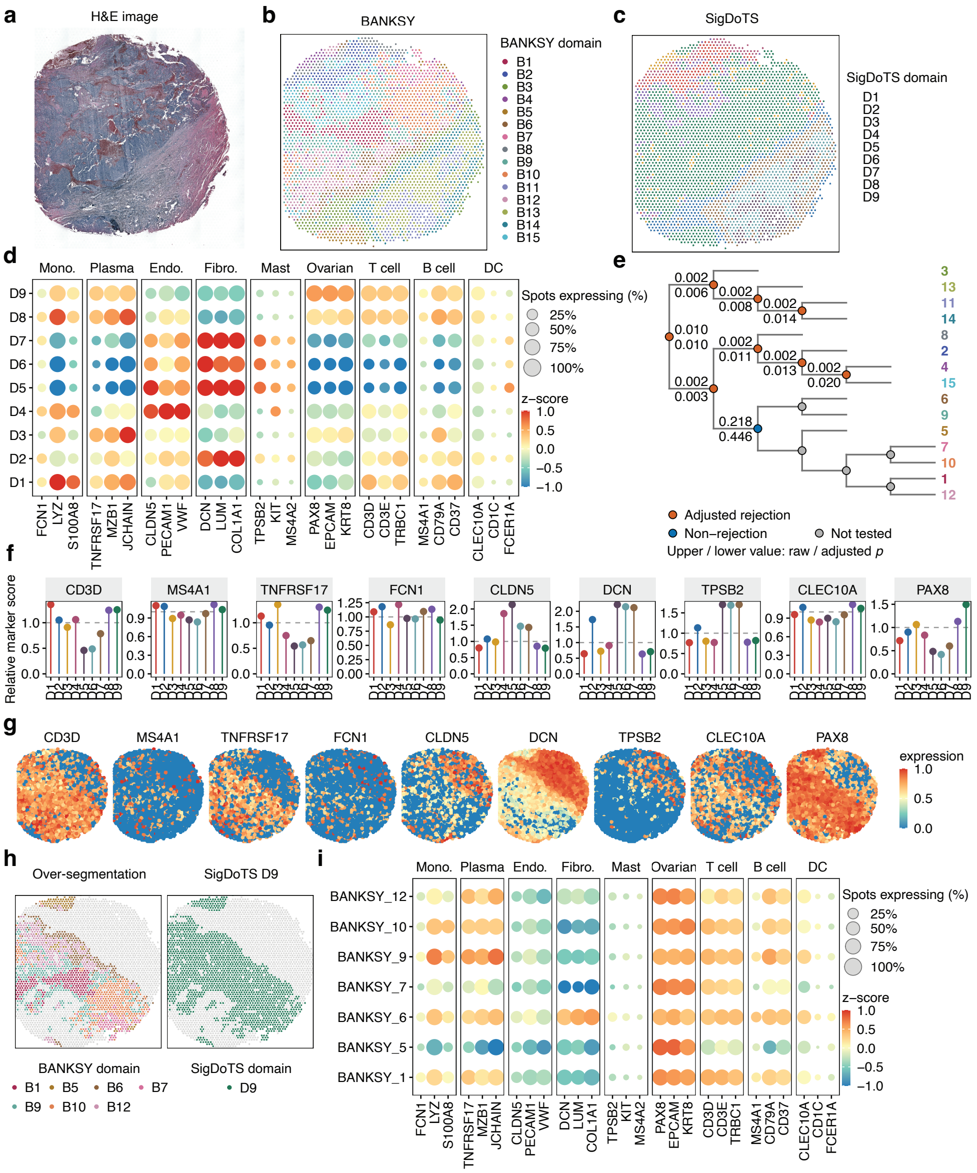

**Supplementary Fig. S6: Nine spatial domains retained by SigDoTS in ovarian cancer Visium data.** **a** H&E-stained ovarian cancer FFPE tissue section. **b** Fifteen initial candidate domains generated by BANKSY (B1–B15). **c** Nine spatial domains retained by SigDoTS after hierarchical testing (D1–D9). **d** Expression of predefined RNA markers representing monocyte, plasma-cell, endothelial, fibroblast, mast-cell, ovarian cancer epithelial, T-cell, B-cell and dendritic-cell programs across the retained domains. Dot color denotes standardized expression ($z$-score), and dot size denotes the percentage of spots expressing the corresponding gene. **e** SigDoTS hierarchical testing tree. Numbers beside the nodes indicate unadjusted $p$-values and hierarchically adjusted $p$-values, and node color indicates rejection or failure to reject after hierarchical adjustment, or that the node was not tested. **f** Relative marker scores of representative RNA markers within each retained domain. **g** Spatial distributions of the corresponding RNA markers within the tissue section after marker-wise normalization. **h** Retained domain D9 and its corresponding BANKSY candidate domains B1, B5, B6, B7, B9, B10 and B12, showing the spatial correspondence between the BANKSY candidate domains and retained domain D9. **i** Expression of predefined RNA markers across the BANKSY candidate domains within D9. Dot color denotes standardized expression ($z$-score), and dot size denotes the percentage of spots expressing the corresponding gene.

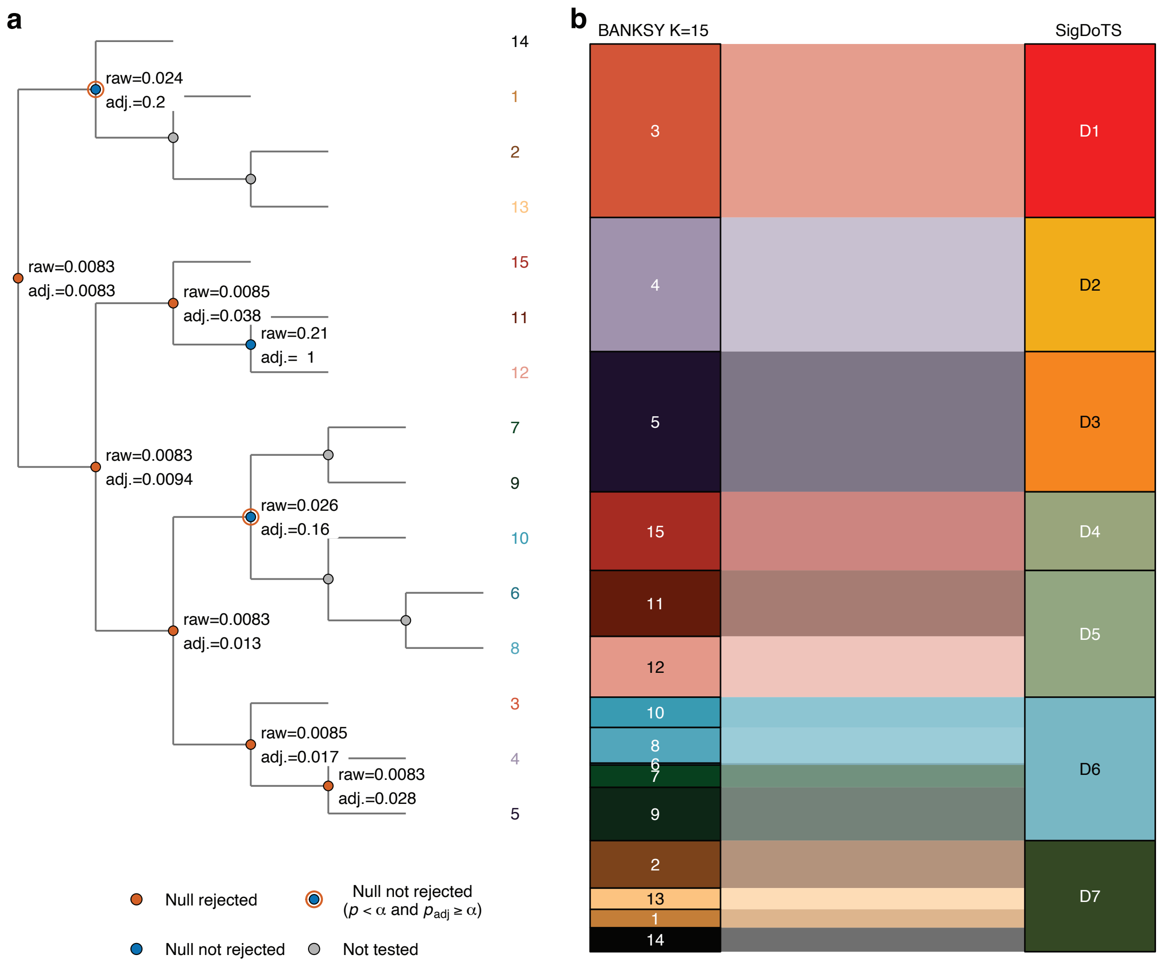

**Supplementary Fig. S7: SigDoTS retains seven spatial domains from 15 BANKSY candidate domains in colorectal adenocarcinoma. a** Hierarchical testing tree for the 15 BANKSY candidate domains. Labels at tested nodes give the unadjusted $p$-value and the hierarchically adjusted $p_{\text{adj}}$-value. Red nodes indicate rejection of the single-domain null after hierarchical adjustment; blue nodes indicate failure to reject before adjustment; red-outlined blue nodes indicate $p < \alpha$ before adjustment but $p_{\text{adj}} \geq \alpha$ after adjustment; and gray nodes were not tested under the hierarchical stopping rule. Leaf-label colors denote the retained SigDoTS domains. **b** Alluvial mapping of the 15 BANKSY candidate domains ($K_0 = 15$) to the seven retained SigDoTS domains, D1–D7. Block heights and band widths represent the numbers of cells assigned to each candidate and retained domain.

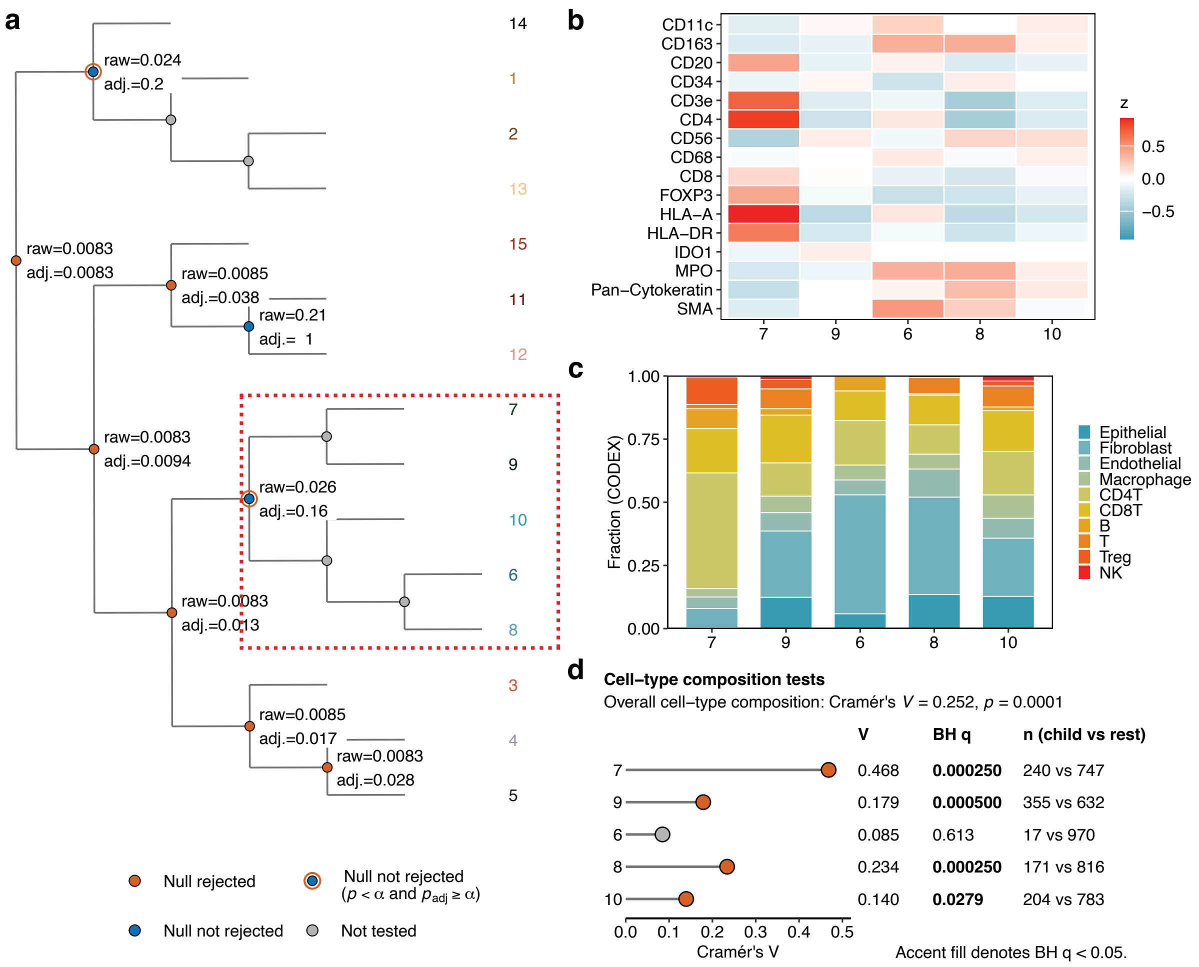

**Supplementary Fig. S8: CODEX cell-type composition differs among candidate domains merged into D6 in the colorectal adenocarcinoma Xenium dataset. a** Hierarchical testing tree with the D6 candidate split outlined in red. **b** Mean CODEX protein intensities, displayed as marker-wise $z$-scores, for adjacent-section cells assigned to candidate domains 7, 9, 6, 8 and 10. **c** CODEX cell-type fractions within the five candidate domains. **d** Overall and candidate-domain-versus-rest analyses of CODEX cell-type composition. Points show Cramér's $V$ for each candidate domain compared with the remaining candidate domains pooled. The table reports Benjamini–Hochberg (BH)-adjusted $q$ values and the corresponding CODEX cell counts. Accent-filled points denote BH-adjusted $q < 0.05$. CODEX measurements were obtained from the adjacent section and were not used for candidate-domain detection or hierarchical testing.

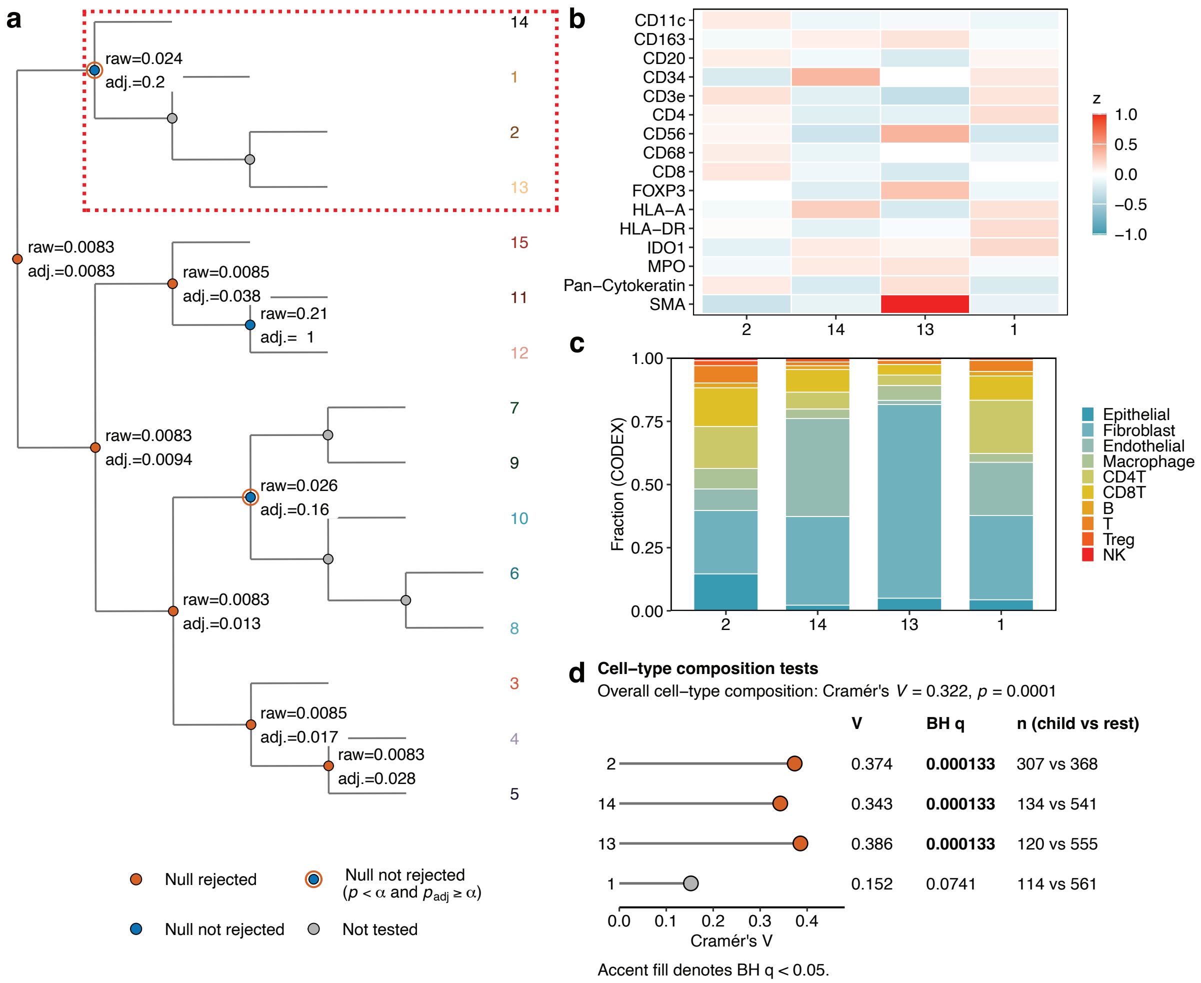

**Supplementary Fig. S9: CODEX cell-type composition differs among candidate domains merged into D7 in the colorectal adenocarcinoma Xenium dataset.** **a** Hierarchical testing tree with the D7 candidate split outlined in red. **b** Mean CODEX protein intensities, displayed as marker-wise $z$-scores, for adjacent-section cells assigned to candidate domains 2, 14, 13 and 1. **c** CODEX cell-type fractions within the four candidate domains. **d** Overall and candidate-domain-versus-rest analyses of CODEX cell-type composition. Points show Cramér's $V$ for each candidate domain compared with the remaining candidate domains pooled. The table reports BH-adjusted $q$ values and the corresponding CODEX cell counts. Accent-filled points denote BH-adjusted $q < 0.05$. CODEX measurements were obtained from the adjacent section and were not used for candidate-domain detection or hierarchical testing.

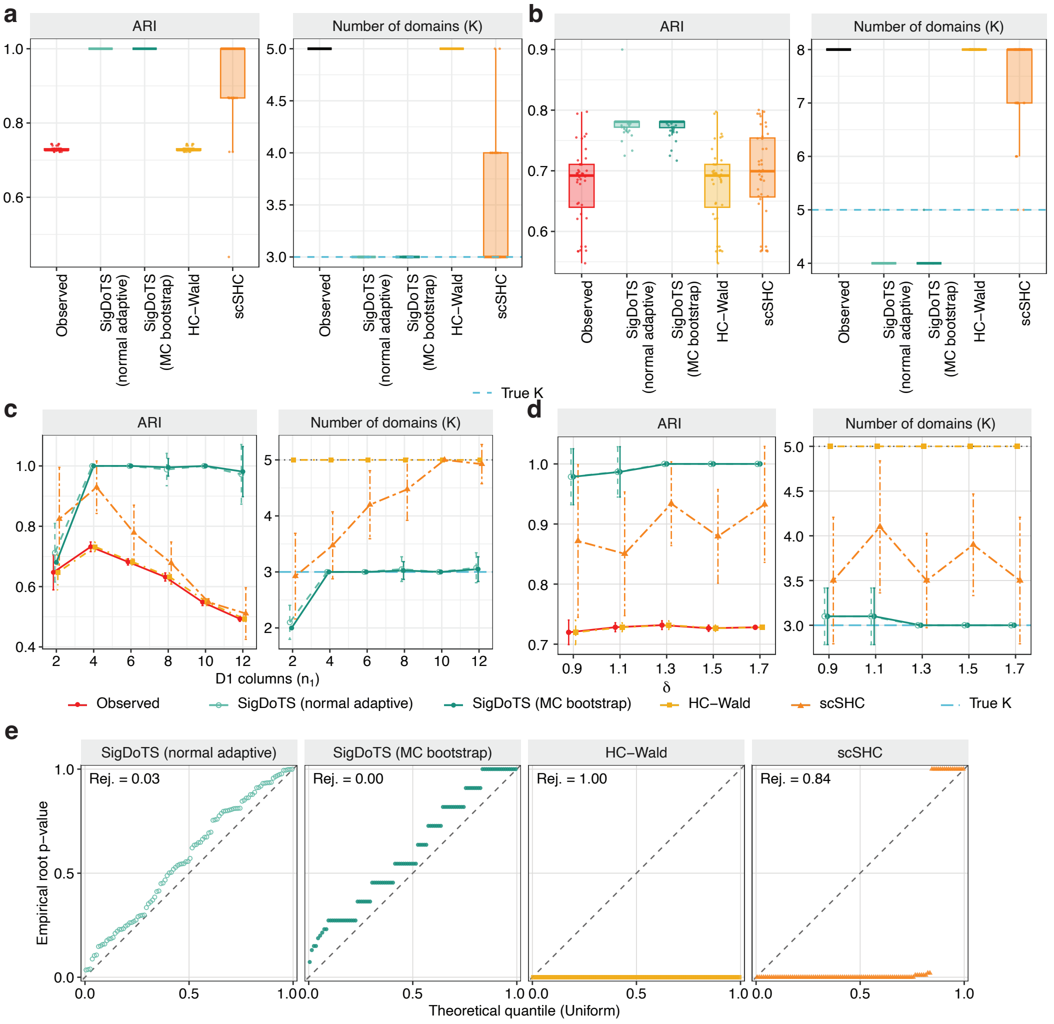

**Supplementary Fig. S10: Comparison of normal-adaptive and Monte Carlo bootstrap implementations of SigDoTS in simulations with a baseline size of 144 spatial units. a** Adjusted Rand index (ARI; left) and number of retained domains $K$ (right) for the three-domain alternative on a $12 \times 12$ grid ($K_{\text{true}} = 3$, initial candidate-domain count $K_0 = 5$; 40 replicates). "Observed" denotes the initial candidate domains. The blue dashed line marks $K_{\text{true}}$; the black bar marks $K_0$. Points denote individual datasets; boxplots show medians and interquartile ranges, with whiskers reaching the most extreme values within 1.5× the interquartile range of the box edges. **b** ARI and retained $K$ for the five-domain alternative on a $20 \times 20$ grid ($K_{\text{true}} = 5$, $K_0 = 8$; 40 replicates). **c** ARI and retained $K$ as domain D1 varied from $n_1 = 2$ to 12 columns, with four columns in each remaining domain and 12 rows ($\delta = 1.3$; 40 replicates per setting). Points and lines show means; error bars indicate ±1 s.d. Blue dashed and gray dotted lines mark $K_{\text{true}} = 3$ and $K_0 = 5$, respectively. **d** ARI and retained $K$ as the mean-difference parameter $\delta$ varied from 0.9 to 1.7 on a $12 \times 12$ grid (10 replicates per setting), with plotting conventions as in **c**. Larger $\delta$ indicates greater differences between domain means. **e** Root $p$-value quantile–quantile plots for 100 global-null datasets on a $12 \times 12$ grid. "Rej." denotes the root rejection rate at $\alpha = 0.05$; diagonals mark the uniform reference.

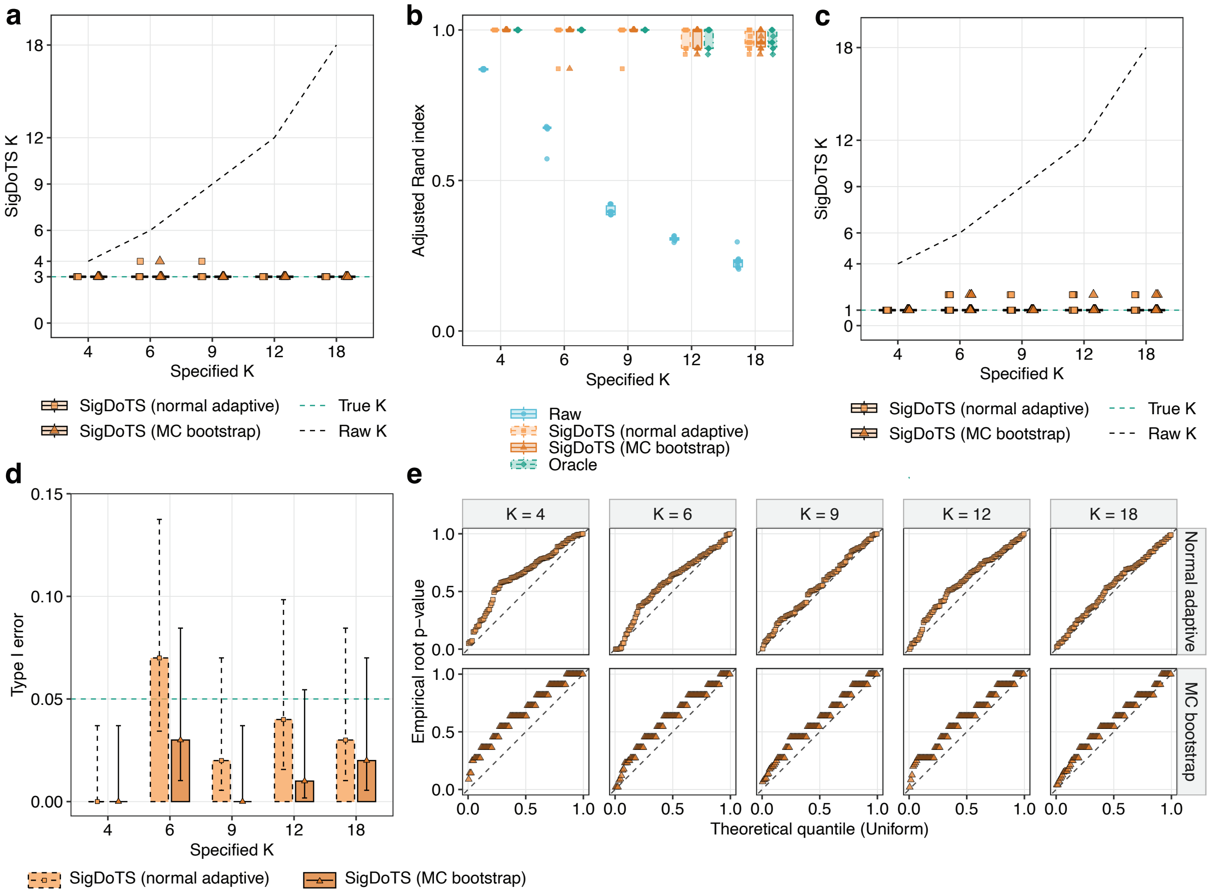

**Supplementary Fig. S11: Comparison of normal-adaptive and Monte Carlo bootstrap implementations of SigDoTS across initial candidate-domain counts.** $k$-means generated $K_0 \in \{4, 6, 9, 12, 18\}$ initial candidate domains, with 100 global-null and 10 three-domain alternative datasets per setting ($\delta = 1.3$). **a** Number of retained domains under the alternative. Reference lines mark $K_{\text{true}} = 3$ and $K_0$. Points denote individual datasets; boxplots show medians and interquartile ranges, with whiskers reaching the most extreme values within 1.5× the interquartile range of the box edges. **b** Adjusted Rand index (ARI) under the alternative, with points and boxplots defined as in **a**. "Raw" denotes the initial candidate domains; "Oracle" denotes merging initial candidate domains along the hierarchy to maximize ARI against the true domain labels, providing an upper bound on domain recovery achievable within that hierarchy. **c** Number of retained domains under the global null, with points and boxplots defined as in **a**. Reference lines mark $K_{\text{true}} = 1$ and $K_0$. **d** Empirical type I error under the global null. Bars show the proportion of datasets with root rejection, error bars show two-sided 95% Wilson confidence intervals and the dashed line marks $\alpha = 0.05$. **e** Root $p$-value quantile–quantile plots under the global null; diagonals mark the uniform reference.

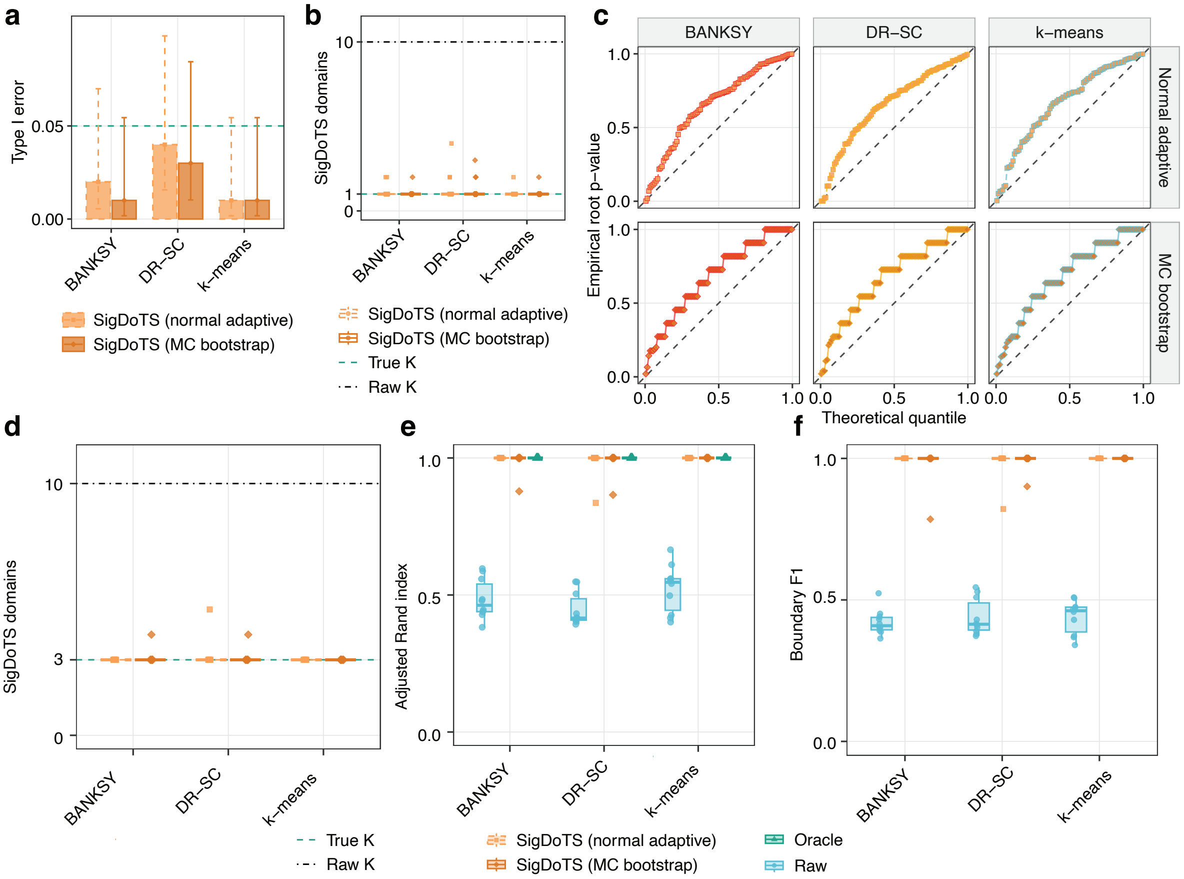

**Supplementary Fig. S12: Comparison of normal-adaptive and Monte Carlo bootstrap implementations of SigDoTS across upstream methods.** BANKSY, DR-SC and $k$-means generated $K_0 = 10$ initial candidate domains, with 100 global-null and 10 three-domain alternative datasets per method ($\delta = 1.3$). **a** Empirical type I error under the global null. Bars show the proportion of datasets in which the root test rejected the single-domain null, error bars show two-sided 95% Wilson confidence intervals and the dashed line marks $\alpha = 0.05$. **b** Number of retained domains under the global null. Reference lines mark $K_{\text{true}} = 1$ and $K_0 = 10$. Points denote individual datasets; boxplots show medians and interquartile ranges, with whiskers reaching the most extreme values within 1.5× the interquartile range of the box edges. **c** Root $p$-value quantile–quantile plots under the global null; diagonals mark the uniform reference. **d** Number of retained domains under the alternative. Reference lines mark $K_{\text{true}} = 3$ and $K_0 = 10$. **e** Adjusted Rand index (ARI) under the alternative. “Raw” denotes the initial candidate domains; “Oracle” denotes merging initial candidate domains along the hierarchy to maximize ARI against the true domain labels, providing an upper bound on domain recovery achievable within that hierarchy. **f** Boundary F1 under the alternative, comparing the initial candidate domains (“Raw”) with domains retained by SigDoTS.
